# BRWD1 establishes epigenetic states for germinal center initiation, maintenance, and function

**DOI:** 10.1101/2024.04.25.591154

**Authors:** Nathaniel E. Wright, Domenick E. Kennedy, Junting Ai, Margaret L. Veselits, Mary Attaway, Young me Yoon, Madeleine S. Durkee, Jacob Veselits, Mark Maienschein-Cline, Malay Mandal, Marcus R. Clark

## Abstract

Germinal center (GC) B cells segregate into three subsets that compartmentalize the antagonistic molecular programs of selection, proliferation, and somatic hypermutation. In bone marrow, the epigenetic reader BRWD1 orchestrates and insulates the sequential stages of cell proliferation and *Igk* recombination. We hypothesized BRWD1 might play similar insulative roles in the periphery. In *Brwd1*^-/-^ follicular B cells, GC initiation and class switch recombination following immunization were inhibited. In contrast, in *Brwd1*^-/-^ GC B cells there was admixing of chromatin accessibility across GC subsets and transcriptional dysregulation including induction of inflammatory pathways. This global molecular GC dysregulation was associated with specific defects in proliferation, affinity maturation, and tolerance. These data suggest that GC subset identity is required for some but not all GC-attributed functions. Furthermore, these data demonstrate a central role for BRWD1 in orchestrating epigenetic transitions at multiple steps along B cell developmental and activation pathways.

## INTRODUCTION

Central to initiation of adaptive humoral immunity is the activation and differentiation of B cells located within the follicles of secondary lymphoid organs^1^. In T cell-dependent responses, activated follicular (Fol) B cells may differentiate into germinal center (GC) B cells over four days, a multi-stage process which includes upregulation of the transcription factor (TF) BCL6^2–7^. This pathway from Fol B cell to GC B cell involves large shifts in molecular programs, including changes in TF networks, chromatin accessibility, and three-dimensional (3D) chromatin architecture^8–12^. It is during this transition from Fol to GC B cells that B cells undergo class switch recombination (CSR)^13^.

Following GC entry, B cells further differentiate. While histologically the GC has two zones, a dark zone (DZ) and a light zone (LZ), we have demonstrated that the DZ contains two distinct populations of B cells that differ in their location and molecular programs^14^. This three-zone model organizes fundamentally different GC programs into specific cell states^15,16^. First, in the LZ, GC B cells are selected by capturing antigen displayed on follicular dendritic cells and presenting this antigen to T follicular helper cells (Tfh) cells^17–27^. Second, in the DZ proliferation (DZp), B cells undergo mitosis or apoptosis adjacent to tingible body macrophages. Third, in the DZ differentiation (DZd), B cells exit the cell cycle and upregulate the mechanisms of somatic hypermutation (SHM). B cells rapidly cycle through the GC until they differentiate into memory B cells (MBCs) or plasma cells (PCs)^1^. Consistent with their functional specialization, the three GC B cell subsets have large differences in transcriptional programs and chromatin accessibility^14^. However, the mechanisms controlling rapid transit through these three very different GC B cell states is unknown.

B cell progenitors also experience large shifts in molecular programs as they develop in the bone marrow^28,29^. Specifically, the epigenetic reader bromodomain and WD repeat-containing protein 1 (BRWD1) mediates the transition between proliferating large pre-B cells and small pre-B cells undergoing light chain recombination via multiple mechanisms^30–32^. At the local scale, BRWD1 repositions nucleosomes relative to DNA GAGA motifs to open and close chromatin^30^. BRWD1 also mediates long-range chromatin contacts over a megabase scale by converting static to dynamic cohesin capable of loop extrusion^32^. In these ways, BRWD1 coordinates the transcriptional, epigenetic, and chromatin 3D transitions necessary for B cell progenitors to stop proliferation and begin recombination. BRWD1 is also highly expressed in both Fol and GC B cells, yet its contributions to the cell state transitions between each population in response to antigen have not been explored.

Herein, we investigate the role of BRWD1 in peripheral B cells using *Brwd1*-floxed mice to delete *Brwd1* in Fol and GC B cells. We demonstrate that BRWD1 is important for GC initiation and CSR in Fol B cells. In GC B cells, we find that loss of *Brwd1* results in a striking collapse of epigenetic transitions and dysregulation of transcriptional programs within the three GC B cell subsets. These results and previous studies^30–32^ indicate that at multiple stages, BRWD1 orchestrates B cell transitions between proliferation and states in which DNA is recombined or mutated to generate diversity and shape immune responses to infection.

## RESULTS

### Design of *Brwd1*-floxed mouse

*Brwd1* is first induced in small pre-B cells during B lymphopoiesis in the bone marrow where it mediates widespread epigenetic changes^30,31^. Using RNA-sequencing (RNA-seq) from different B cell populations in WT mice, we found that *Brwd1* expression was over twice as high in Fol B cells than in small pre-B cells (Extended Data Fig 1a). Furthermore, *Brwd1* expression changed across GC B cell subsets, with the greatest expression in the DZd^14^. This dynamic regulation of *Brwd1* in Fol and GC B cells suggested that BRWD1 may mediate both transitions between and maintenance of peripheral B cells.

To study the role of BRWD1 in peripheral C57BL/6 B cells, we derived a *Brwd1*-floxed allele with Lox71 and Lox66 sites in opposite orientations surrounding *Brwd1* exons 6, 7, and 8 (Extended Data Fig 1b). We first crossed these mice with mice expressing *Cre* under the *Fcer2a* promoter (*Cd23^Cre^*)^33^. The targeting construct includes an inverse tdTomato (tdT) reporter so that following Cre recombination, exons 6, 7, and 8 are inverted and tdT is expressed (Extended Data Fig 1c). Using RNA-seq, we confirmed that tdT^+^ Fol B cells in *Cd23^Cre/wt^ Brwd1^fl/fl^* (*Brwd1*-KO^Fol^) mice do not transcribe exons 6, 7, and 8 of *Brwd1* (Extended Data Fig 1d). Homozygous *Brwd1*-KO^Fol^ mice had greater tdT median fluorescence intensity (MFI) than heterozygous *Cd23^Cre/wt^ Brwd1^fl/wt^* (*Brwd1*-Het^Fol^) mice, demonstrating that both alleles were deleted in homozygous mice (Extended Data Fig 1e).

### BRWD1 is not necessary for maintenance of Fol B cells

To study whether BRWD1 is important for the maintenance of peripheral B cells, we characterized *Brwd1*-KO^Fol^ mice at the steady state. Flow cytometry demonstrated that control (*Brwd1^fl/fl^*), *Brwd1*-Het^Fol^, and *Brwd1*-KO^Fol^ mice had comparable numbers of CD20^+^CD19^+^CD93^-^ mature B cells in the spleen (Extended Data Fig 2a-b). Additionally, these mice had the same number and frequency of Fol and marginal zone (MZ) B cells (Extended Data Fig 2c-f). Approximately 96% of Fol B cells and 82% of MZ B cells expressed tdT, confirming efficient Cre recombination in *Brwd1*-KO^Fol^ mice (Extended Data Fig 2g-h). Together, these results demonstrate that steady state splenic B cells do not require BRWD1.

### BRWD1 is important for GC initiation

Upon antigenic stimulation, Fol B cells differentiate into GC B cells. To determine whether BRWD1 is important for the differentiation of Fol B cells, we intraperitoneally immunized *Brwd1*-KO^Fol^ mice with sheep red blood cells at days 0 and 5 and analyzed spleens 14 days post-immunization at the peak of the GC response (Fig 1a-b)^34^. The number of B220^+^ B cells was equivalent between *Brwd1*-KO^Fol^ and control mice (Fig 1c). However, the number and frequency of GL7^+^CD95^+^ GC B cells were decreased over twofold in the *Brwd1*-KO^Fol^ mice (Fig 1d-e). This decrease was observed in GC B cells within the LZ, DZp, and DZd (Fig 1f). Over 85% of GC B cells expressed tdT in the *Brwd1*-KO^Fol^ mice, demonstrating that the smaller GC population primarily contained *Brwd1*^-/-^ GC B cells (Fig 1g). *Brwd1*-KO^Fol^ mice had no significant change in the number of Tfh cells but did have a decreased Tfh cell frequency (Fig 1h-i)^35^. Furthermore, the frequency of GC B cells and Tfh cells positively correlated with one another, showing that the decreased Tfh cell frequency was consistent with the smaller GC B cell response (Fig 1j)^35^.

**Figure 1.**
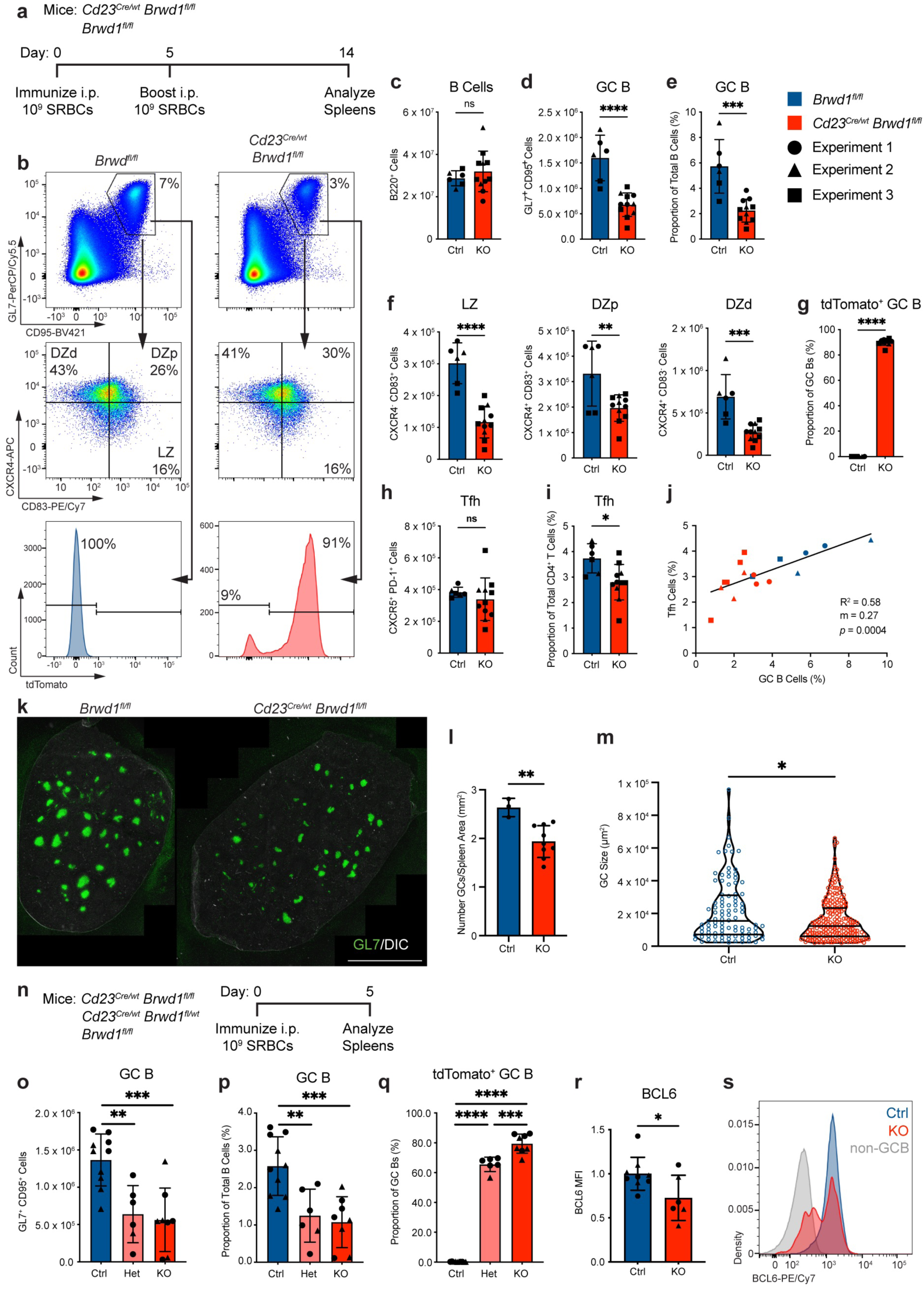
Deletion of *Brwd1* in Fol B cells represses GC initiation. **(a)** *Cd23^Cre/wt^ Brwd1^fl/fl^* (KO) and *Brwd1^fl/fl^* (Ctrl) mice were immunized intraperitoneally (i.p.) with 10^9^ sheep red blood cells (SRBCs) at days 0 and 5. Spleens were collected at day 14 post-immunization (Ctrl n = 6 mice, KO n = 11 mice). **(b)** Representative flow plots of GC B cells first gated on live B220^+^ cells. **(c)** Total number of B220^+^ B cells. **(d-e)** Absolute number (d) and frequency (e) of GL7^+^CD95^+^ GC B cells. **(f)** Number of LZ, DZp, and DZd GC B cells. **(g)** Proportion of GC B cells expressing tdTomato. **(h-i)** Total number (h) and frequency (i) of CXCR5^+^PD-1^+^ T follicular helper (Tfh) cells. **(j)** Linear regression of frequency of GC B cells vs. frequency of Tfh cells. m = slope. **(k)** Representative images of spleen sections from *Cd23^Cre/wt^ Brwd1^fl/fl^* (KO) and *Brwd1^fl/fl^* (Ctrl) mice imaged at 10x magnification and stained with GL7. (DIC = differential interference contrast, Scale bar = 2,000 µm, KO n = 3 mice, Ctrl n = 9 mice) **(l)** Number of GCs per spleen area. **(m)** Size of GCs as measured by area of GL7 staining. Violin plots show median and quartiles (Ctrl n = 10 mice, Het n = 6 mice, KO n = 8 mice). **(n)** *Cd23^Cre/wt^ Brwd1^fl/fl^* (KO), *Cd23^Cre/wt^ Brwd1^fl/wt^* (Het), and *Brwd1^fl/fl^* (Ctrl) mice were immunized i.p. with 10^9^ SRBCs, and spleens were harvested at day 5. **(o-p)** Total number (o) and frequency (p) of GL7^+^CD95^+^ GC B cells. **(q)** Proportion of GC B cells expressing tdT. **(r)** Median fluorescence intensity (MFI) of BCL6 in GC B cells. Control data was normalized to 1 to account for voltage differences between flow cytometry experiments (Ctrl n = 9 mice, KO n = 6). **(s)** Representative histogram of BCL6 intensity in Ctrl and KO GC B cells and in non-GC B220^+^CD95^-^GL7^-^ B cells. (\**p* < 0.05, \*\**p* < 0.01, \*\*\**p* < 0.001, \*\*\*\**p* < 0.0001, m: two-sided Mann-Whitney, j: simple linear regression, all others: two-sided unpaired *t*-test, bar plots show mean ± standard deviation)

We confirmed a defect in the GC response by immunofluorescence microscopy. Spleens from *Brwd1*-KO^Fol^ mice had fewer GCs per splenic area than controls (Fig 1k-l). Additionally, GCs from *Brwd1*-KO^Fol^ mice were smaller compared to controls as measured by GL7 fluorescence (Fig 1m). Together, these results demonstrate that BRWD1 is important for a proper GC response.

This diminished GC B cell response in *Brwd1*-KO^Fol^ mice could reflect either a defect in initial differentiation of Fol B cells into GC B cells or in expansion and maintenance of GC B cells. Because GCs form around day 4 post-immunization, we measured the GC response at day 5 post-immunization (Fig 1n)^3,5^. At this time point, we observed a 2.5-fold decrease in both the number and frequency of GC B cells, suggesting that BRWD1 is important for GC initiation (Fig 1o-p). Approximately 80% of GC B cells expressed tdT, suggesting that *Brwd1*^-/-^ GC B cells can populate the GC once the response is initiated (Fig 1q).

We considered and ruled out several possibilities for this defect in GC initiation. First, the decreased GC response in *Brwd1*-KO^Fol^ mice at day 5 post-immunization was not caused by *Brwd1*^-/-^ B cells alternatively differentiating into plasmablasts, because there was no significant difference in the number or frequency of CD138^+^ plasmablasts compared to controls (Extended Data Fig 3a-c). Next, GC B cells in *Brwd1*-KO^Fol^ mice had no difference in the prevalence of apoptosis or cell death as measured by Annexin V or fluorescent inhibitors of caspase (FLICA) (Extended Data Fig 3d-g). The proliferating subpopulation among GC B cells is marked by the highest CD83 and CXCR4 expression^14^. There was no difference in the proportion of proliferating cells among the CD83^high^CXCR4^high^ population as measured by DAPI^high^ cells in the S, G2, or M phases of the cell cycle (Extended Data Fig 3h). Furthermore, there was no change in the DZp to LZ ratio or the DZd to LZ ratio (Extended Data Fig 3i-j).

We next examined the expression of BCL6 because of its role in GC differentiation. GC B cells from *Brwd1*-KO^Fol^ mice expressed lower levels of BCL6 as measured by flow cytometry (Fig 1r). This was due to a bimodal BCL6 distribution where some GC B cells had intermediate BCL6 expression between that of control GC B cells and naïve B220^+^CD95^-^GL7^-^ B cells (Fig 1s). The GC precursor population expresses intermediate BCL6 and peaks around day 4 post-immunization^5^. Thus, these results further support that BRWD1 is necessary for the initial differentiation of Fol B cells into GC B cells.

### BRWD1 facilitates class switch recombination

CSR requires changes in chromatin topology at the class switch locus^36^. Therefore, we investigated whether BRWD1 is important for CSR. Indeed, at day 14 post-immunization, fewer tdT^+^ GC B cells were IgM^-^ when compared with tdT^-^ cells from the same mouse, suggesting that *Brwd1*^-/-^ B cells undergo less CSR (Fig 2a). Next, we cultured Fol B cells *in vitro* with IL-4 and TGFβ to induce class switching to IgG1. tdT^+^ cells from *Brwd1*-KO^Fol^ mice had decreased switching to IgG1 compared to controls (Fig 2b-c). tdT^-^ cells from *Brwd1*-KO^Fol^ mice had no change in IgG1 class switching, demonstrating that *Brwd1* plays a cell-intrinsic role in CSR.

**Figure 2.**
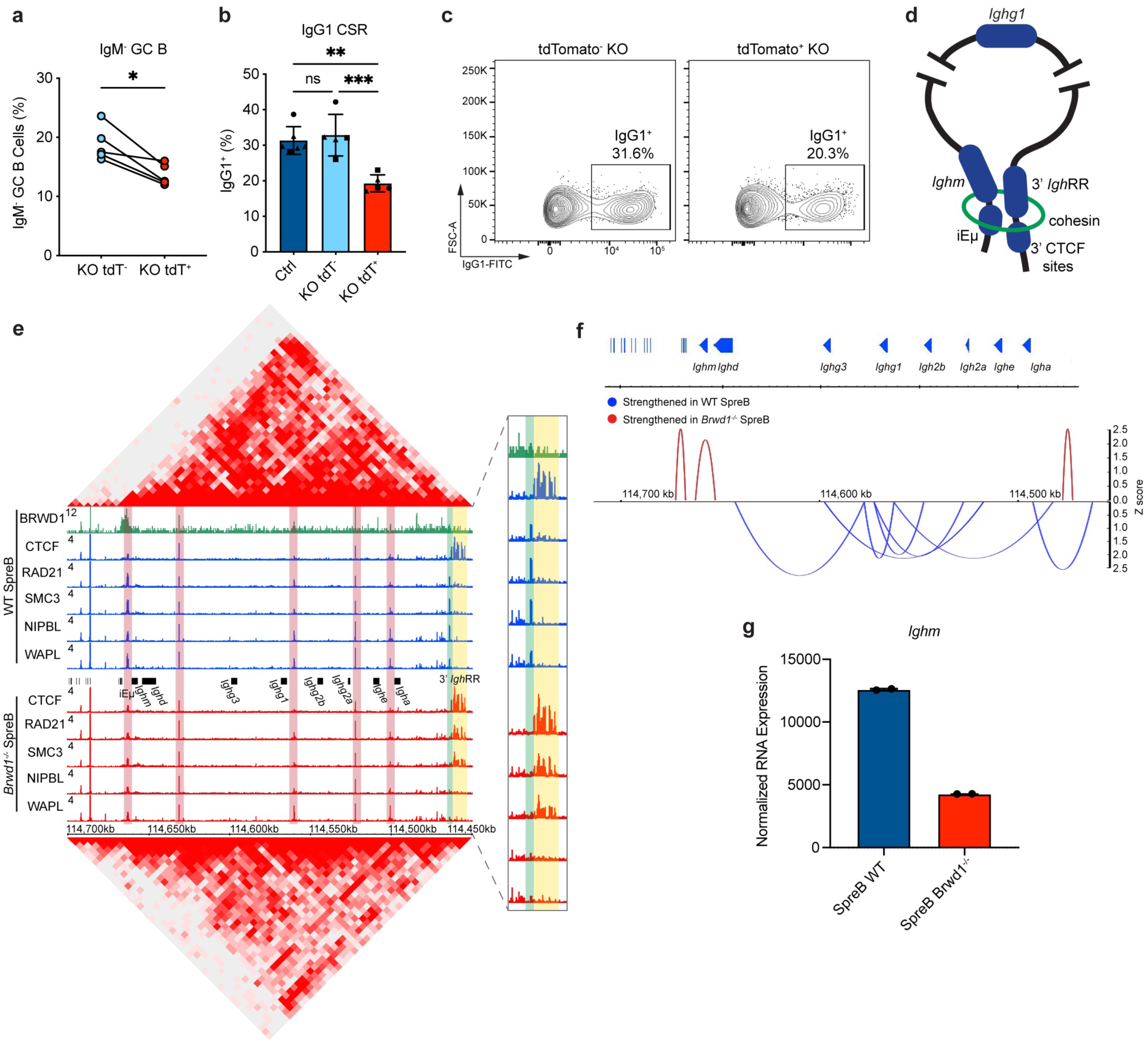
BRWD1 enhances class switch recombination. **(a)** Frequency of IgM^-^ cells after gating on tdT^-^ or tdT^+^ GC B cells. *Cd23^Cre/wt^ Brwd1^fl/fl^* (KO) mice were immunized intraperitoneally with sheep red blood cells at days 0 and 5. Spleens were analyzed day 14 post-immunization (KO n = 5 mice). **(b)** *In vitro* class switching to IgG1. Splenic B cells from KO and *Brwd1^fl/fl^* (Ctrl) mice were cultured for 96 hours with LPS and IL-4, and class switching was measured by flow cytometry (Ctrl n = 6 mice, KO n = 5 mice). **(c)** Representative flow plots of IgG1^+^ frequency in tdT^-^ or tdT^+^ cells from KO mice. First gated on CD19^+^ cells. **(d)** Model of chromatin looping between the 5’ iEµ enhancer and the 3’ *Igh* regulatory region (*Igh*RR) at the class switch locus. The full cohesin ring complex (green) mediates loop extrusion. **(e)** Hi-C heatmap, ChIP-seq tracks for BRWD1, CTCF, RAD21, SMC3, NIPBL, and WAPL in WT (top) and *Brwd1^-/-^* (bottom) small pre-B cells (SpreB). Analysis of data sets previously published^32^. The dynamic cohesin complex is highlighted green, the static cohesin complex is highlighted yellow, and invariant cohesin complexes at CTCF sites are highlighted red. ChIP-seq tracks showing the dynamic and static cohesin complexes are magnified. Data is representative of n = 2 data sets. **(f)** Arc plots indicating strengthened Hi-C interactions (*p* < 0.05) in WT or *Brwd1^-/-^* small pre-B cells across the class switch locus at 5 kb resolution. **(g)** RNA-seq expression of *Ighm* in WT and *Brwd1^-/-^* small pre-B cells (n = 2 per group). (\**p* < 0.05, \*\**p* < 0.01, \*\*\**p* < 0.001, two-sided unpaired *t*-test, bar plots show mean ± standard deviation)

Class switching requires precise chromatin loop extrusion across the class switch locus^36,37^. During B cell development, a loop across the entire locus is established that brings together the 5’ iEµ enhancer and the 3’ *Igh* regulatory region (Fig 2d), and this loop is maintained in naïve Fol B cells^37,38^. Because BRWD1 regulates cohesin to modulate chromatin looping, we measured BRWD1 and components of the cohesin complex at the class switch locus^32^. Chromatin immunoprecipitation followed by sequencing (ChIP-seq) revealed that in small pre-B cells, BRWD1 binds at multiple sites within the *Igh* locus, with an especially strong peak at iEµ (Fig 2e). In WT cells, we observed a dynamic cohesin complex containing coincident NIPBL and WAPL at the 3’ end of the locus (Fig 2e, highlighted green)^32^. In *Brwd1*^-/-^ small pre-B cells, the 3’ dynamic cohesin complex was lost and instead static cohesin was found at the 3’ CTCF sites (highlighted yellow). Invariant cohesin complexes at CTCF sites (highlighted red) were also observed throughout the locus. This loss of the dynamic cohesin complex was associated with decreased Hi-C contacts and decreased looping across the locus (Fig 2e-f). Furthermore, the loss of looping resulted in decreased *Ighm* expression compared to WT small pre-B cells (Fig 2g). Thus, BRWD1 binds at and is necessary for chromatin looping across the class switch locus during B cell development. These data suggest that the BRWD1-dependent chromatin topology established in small pre-B cells is necessary for efficient CSR upon Fol B cell activation.

### BRWD1 establishes epigenetic states for GC initiation

We next studied whether *Brwd1* deletion affects the transcriptional or epigenetic states of resting Fol B cells. We first sorted tdT^+^ from *Brwd1*-KO^Fol^ mice and performed RNA-seq. tdT^+^ Fol B cells clustered with Fol B cells from control mice by principal component analysis (PCA) when compared with WT GC B cells (Fig 3a). Fol B cells were also highly similar by a Pearson correlation (Fig 3b). Only 17 genes (log_2_FC > 1, *q* < 0.05) were differentially expressed between tdT^+^ Fol B cells and control Fol B cells from *Brwd1^fl/fl^* mice (Fig 3c). These results demonstrate that deletion of *Brwd1* in Fol B cells has minimal transcriptional effects.

**Figure 3.**
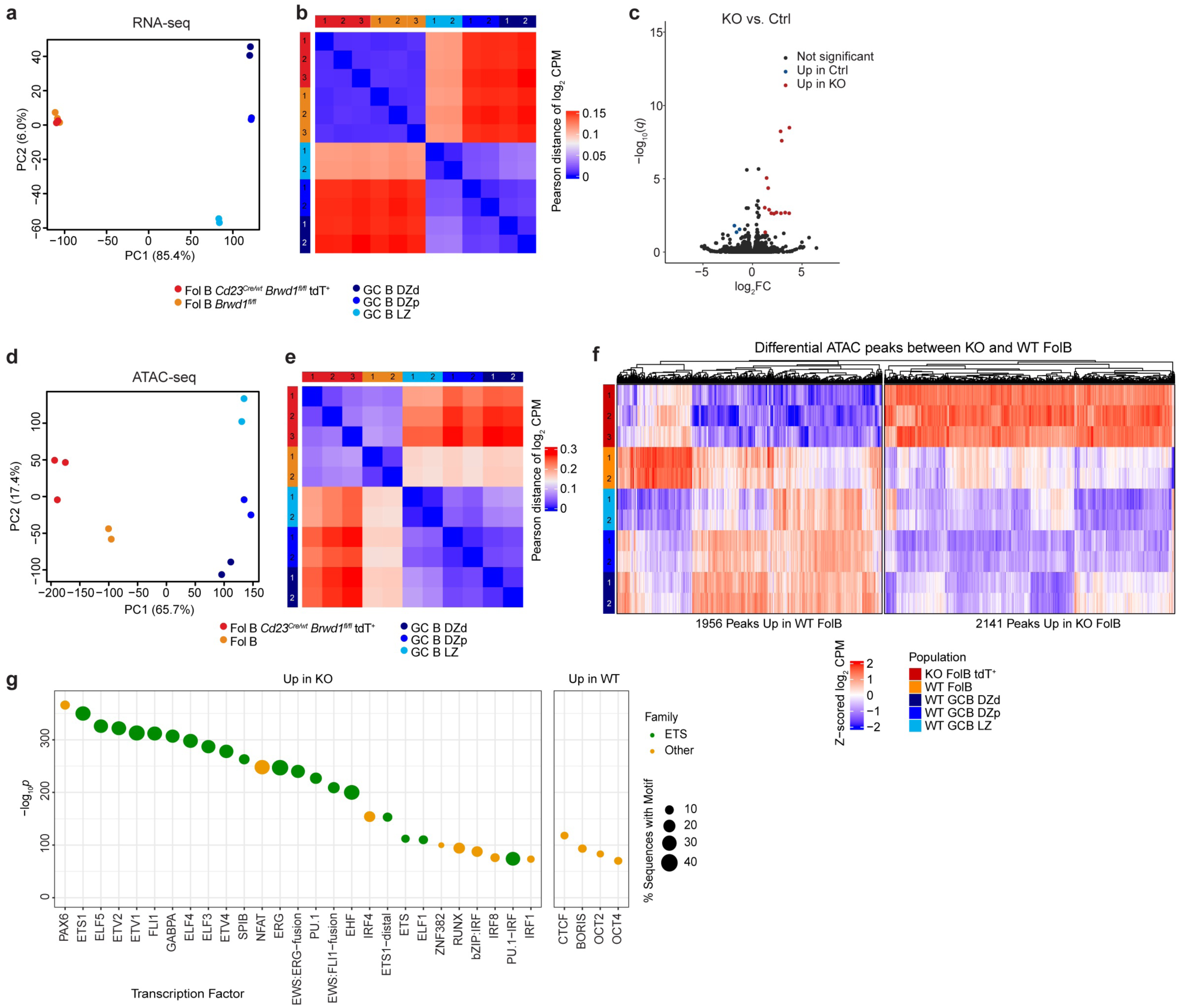
BRWD1 determines genomic accessibility in resting Fol B cells. **(a)** PCA plot of RNA-seq from *Cd23^Cre/wt^ Brwd1^fl/fl^* tdT^+^ Fol B cells (KO), *Brwd1^fl/fl^* Fol B cells (Ctrl), and WT GC B cell subsets (KO Fol B n = 3 mice, Ctrl Fol B n = 3 mice). WT GC B cell RNA-seq (n = 2 for each cell population) was previously published^14^. **(b)** Pearson correlation heatmap of RNA-seq from listed populations. **(c)** Volcano plot of differentially expressed genes (*q* < 0.05, log_2_ fold change > 1) between tdT^+^ *Cd23^Cre/wt^ Brwd1^fl/fl^* (KO) and *Brwd1^fl/fl^* (Ctrl) Fol B cells. **(d)** PCA plot of ATAC-seq from *Cd23^Cre/wt^ Brwd1^fl/fl^* tdT^+^ Fol B cells (KO), WT Fol B cells (Ctrl), and WT GC B cell subsets (KO Fol B n = 3 mice, Ctrl Fol B n = 2 mice). WT GC B cell ATAC-seq (n = 2 for each cell population) was previously published^14^. **(e)** Pearson correlation heatmap of ATAC-seq from listed populations. Note scale differences between b and e. **(f)** Heatmap of differential accessibility peaks (*q* < 0.05, log_2_ fold change > 1) between tdT^+^ *Cd23^Cre/wt^ Brwd1^fl/fl^* and WT Fol B cells. Accessibility by ATAC-seq is shown for listed populations. **(g)** TF motifs enriched in accessible regions up in WT or KO Fol B cells generated using HOMER. TF motifs in the ETS family, which have similar binding sequences, are shown in green, while all other TF motifs are in orange. TF motifs enriched at *p* < 10^-30^ are shown.

To determine whether loss of BRWD1 affects the chromatin accessibility of Fol B cells, we performed ATAC-seq. By PCA, tdT^+^ Fol B cells from *Brwd1*-KO^Fol^ mice clustered separately from both WT Fol B cells and GC B cells (Fig 3d). Chromatin accessibility was more different across Fol B cell populations than the differences observed in transcription (Fig 3e, note scale).

When we compared differential chromatin accessibility peaks (log_2_FC > 1, *q* < 0.05) between tdT^+^ and WT Fol B cells, we observed that 1,956 peaks increased in WT and 2,141 peaks increased in *Brwd1*^-/-^ Fol B cells (Fig 3f). Interestingly, the accessibility of many of these peaks in WT Fol B cells matched the accessibility in GC B cells, suggesting that the accessibility of these sites is maintained by BRWD1 as Fol B cells differentiate into GC B cells. TF motif accessibility was altered between WT and *Brwd1*^-/-^ Fol B cells for TFs important in B cell differentiation (Fig 3g). For example, binding sites for OCT2, which prepares and facilitates Fol to GC B cell differentiation programs, were more accessible in WT Fol B cells^12,39,40^. Conversely, accessibility at Ets family binding sites, which can restrain B cell activation and differentiation programs, were increased in *Brwd1*^-/-^ Fol B cells^41–44^. Accessibility at IRF4 and 8 binding sites, which regulate activated B cell differentiation programs were also increased in *Brwd1*^-/-^ Fol B cells^45,46^. These results suggest that in Fol B cells BRWD1 primes genomic accessibility for transition to the GC state following antigen stimulation.

### BRWD1 restrains GC B cell proliferation

While BRWD1 was important for GC initiation, it is also expressed in GC subsets and therefore may have additional roles once B cells have fully differentiated into GC B cells. To study this, we crossed the *Brwd1*-floxed mice with *Aicda^Cre^* mice to delete *Brwd1* in GC B cells approximately 4 days post-immunization when GCs have begun to form^3,8,47^. We immunized *Aicda^Cre/wt^ Brwd1^fl/fl^* (*Brwd1*-KO^GC^) mice and control *Brwd1^fl/fl^* mice and analyzed spleens after 14 days (Fig 4a-b). We observed no significant difference in the total number of B220^+^ B cells in *Brwd1*-KO^GC^ mice compared with controls (Fig 4c). However, there was a significant increase in the total number and frequency of GC B cells in the *Brwd1*-KO^GC^ mice (Fig 4d-e). In the *Brwd1*-KO^GC^ mice, 80% of GC B cells had deleted *Brwd1* as shown by tdT expression, and this frequency was consistent across GC B cell subsets (Fig 4f, Extended Data Fig 4a). tdT MFI varied across the three GC subsets, with the greatest tdT expression in the LZ (Extended Data Fig 4b-c). The increase in GC B cell number was observed across each of the three GC subsets, with minimal changes in the relative proportion of cells within each GC subset as measured by the DZd/LZ and DZp/LZ ratios (Fig 4g-i). Similar results were observed in heterozygous *Aicda^Cre/wt^ Brwd1^fl/wt^* (*Brwd1*-Het^GC^) mice (Extended Data Fig 4d-g). As a control, *Aicda^Cre/wt^* mice had the same GC B cell frequency as *Brwd1^fl/fl^* mice, demonstrating that the GC defect observed in *Brwd1*-KO^GC^ mice was not due to *Aicda* haploinsufficiency (Extended Data Fig 4h). In total, both *Brwd1* deletion and *Brwd1* haploinsufficiency are sufficient to alter GC B cell numbers.

**Figure 4.**
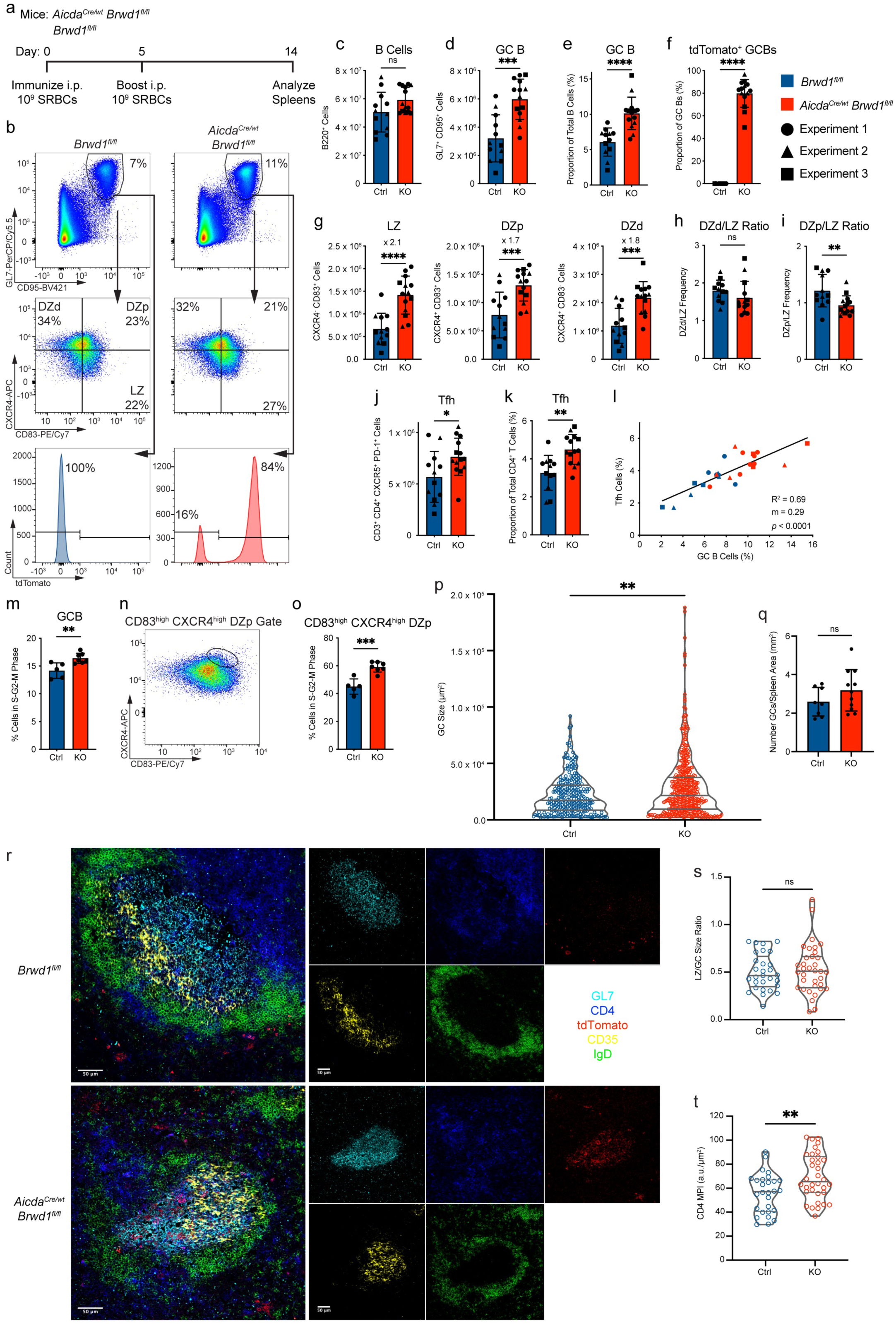
BRWD1 restrains proliferation in GC B cells. **(a)** *Aicda^Cre/wt^ Brwd1^fl/fl^* (KO) and *Brwd1^fl/fl^* (Ctrl) mice were immunized intraperitoneally (i.p.) with 10^9^ sheep red blood cells (SRBCs) at days 0 and 5. Spleens were collected at day 14 post-immunization (Ctrl n = 12 mice, KO n = 14 mice). **(b)** Representative flow plots of GC B cells first gated on B220^+^ cells. **(c)** Total B cell number. **(d-e)** Total number (d) and frequency (e) of GL7^+^CD95^+^ GC B cells. **(f)** Proportion of GC B cells expressing tdT. **(g)** Number of LZ, DZp, and DZd GC B cells. **(h)** Ratio of DZd and LZ frequency. **(i)** Ratio of DZp and LZ frequency. **(j-k)** T follicular helper (Tfh) cell number (j) and frequency (k). **(l)** Correlation of GC B cell and Tfh cell frequency. m = slope. **(m)** Frequency of proliferating GC B cells in the S, G2, or M phases of the cell cycle as measured by DAPI^high^ frequency (Ctrl n = 5 mice, KO n = 7 mice). **(n)** Flow cytometry gate on CXCR4^high^CD83^high^ DZp GC B cells. **(o)** Frequency of proliferating CXCR4^high^CD83^high^ GC B cells. **(p)** GC size measured by GL7 fluorescence (Ctrl n = 8 mice, KO n = 11 mice). **(q)** Number of GCs normalized by spleen area (mm^2^). **(r)** Representative immunofluorescence imaging of spleens at 20x magnification using GL7, anti-CD4, anti-tdT, anti-CD35, and anti-IgD (Ctrl n = 4 mice, 29 GCs; KO n = 4 mice, 35 GCs). **(s)** Ratio of LZ size to total GC size. **(t)** CD4 mean pixel intensity (MPI, arbitrary units/µm^2^) within the LZ per GC imaged. (\**p* < 0.05, \*\**p* < 0.01, \*\*\**p* < 0.001, \*\*\*\**p* < 0.0001, l: simple linear regression, p, s, t: two-sided Mann-Whitney, all others: two-sided unpaired *t*-test, bar plots show mean ± standard deviation, violin plots show median and quartiles)

We next examined the Tfh cell compartment. The number and frequency of CD3^+^CD4^+^CXCR5^+^PD-1^+^ Tfh cells were significantly increased in *Brwd1*-KO^GC^ mice, and the frequency of GC B cells strongly correlated with the frequency of Tfh cells within each spleen (Fig 4j-l, Extended Data Fig 4i). These results are consistent with the increase in GC B cells, because increased antigen presentation to Tfh cells results in greater Tfh cell proliferative expansion^35^. This suggests that antigen presentation in *Brwd1*^-/-^ GC B cells is intact.

To determine whether *Brwd1*^-/-^ GC B cells were undergoing greater proliferation, we stained cells with DAPI to measure the proportion of cells in the S, G2, or M phases of the cell cycle. When gated on total GC B cells, *Brwd1*-KO^GC^ mice had a greater proportion of proliferating GC B cells (Fig 4m). In prior work, we demonstrated that the CXCR4^high^CD83^high^ DZp population within the GC is where GC B cells undergo mitosis, and that expression of both surface markers positively correlates with DNA content^14^. Gating on this CXCR4^high^CD83^high^ DZp population showed an increase in proliferating cells in *Brwd1*-KO^GC^ mice compared to controls (Fig 4n-o, Extended Data Fig 4j). CXCR4^+^CD83^-^ DZd GC B cells also had greater proliferation, suggesting a blending of proliferative programs throughout the GC, although most proliferation was still confined to the DZp (Extended Data Fig 4k-l). Finally, differences in GC B cell numbers were not due to differences in cell viability (Extended Data Fig 4m). These results demonstrate that BRWD1 represses proliferation in GC B cells, consistent with BRWD1’s role as a *Myc* repressor in small pre-B cells^31^.

We further characterized GC responses by immunofluorescence microscopy of spleen sections from *Brwd1*-KO^GC^ mice. This analysis revealed that a portion of GCs in *Brwd1*-KO^GC^ mice were twice as large in cross-sectional area as GCs from control mice, with a greater spread in the distribution of GC size (Fig 4p). There was no significant difference in the number of GCs per spleen area (Fig 4q). GCs from *Brwd1*-KO^GC^ mice had normal architecture with distinct light and dark zones, and there was no difference in the ratio of LZ size to total GC size (Fig 4r-s). In agreement with the Tfh flow cytometry data, there was an increase in the CD4 mean pixel intensity within the LZ (Fig 4t). In summary, *Brwd1* deficiency in GC B cells caused an increase in GC size without affecting GC architecture.

### BRWD1 is not required for post-GC MBCs or PCs

GC B cells can exit GCs by further differentiating into MBCs or PCs. To determine whether BRWD1 is important for these transitions, we studied the prevalence of MBCs and PCs in the spleens of *Brwd1*-KO^GC^ and control *Brwd1^fl/fl^* mice 14 days post-immunization (Fig 5a). We observed no significant difference in the number or frequency of B220^+^CD38^+^GL7^-^IgD^-^ MBCs nor of B220^int^CD138^+^ PCs (Fig 5b-e).

**Figure 5.**
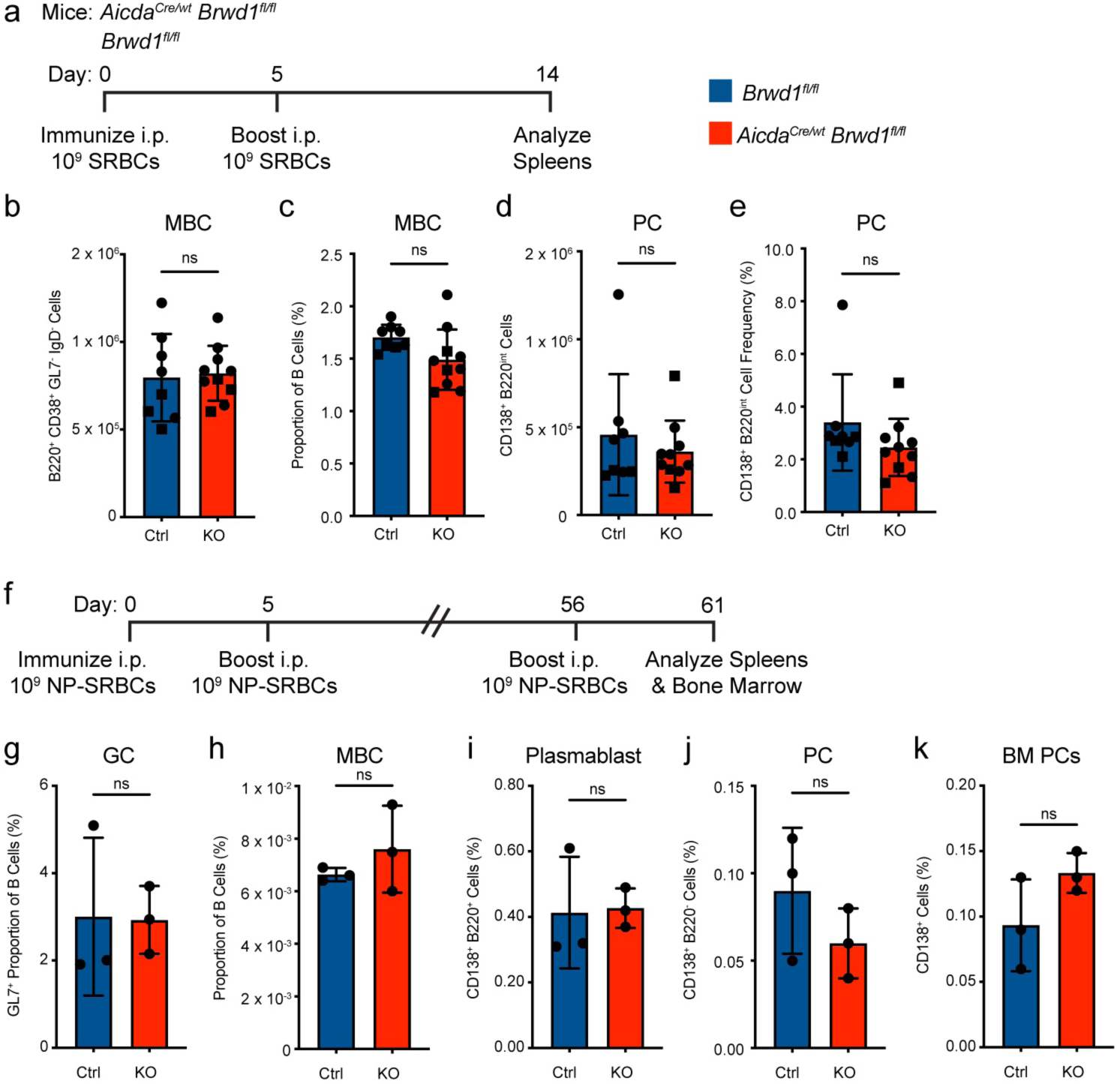
BRWD1 is not required for GC-derived MBCs and PCs. **(a)** *Aicda^Cre/wt^ Brwd1^fl/fl^* (KO) and *Brwd1^fl/fl^* (Ctrl) mice were immunized with SRBCs i.p. and boosted at day 5. Splenic cells were analyzed at day 14 post-immunization (Ctrl n = 8 mice, KO n = 10 mice). **(b)** Number of B220^+^CD38^+^GL7^-^IgD^-^ MBCs. **(c)** Frequency of MBCs as a proportion of all B220^+^ cells. **(d-e)** Number (d) and frequency (e) of CD138^+^B220^int^ PCs. **(f)** *Aicda^Cre/wt^ Brwd1^fl/fl^* (KO) and *Brwd1^fl/fl^* (Ctrl) mice were immunized with NP-SRBCs i.p. at days 0, 5, and 56. Spleens and bone marrow were analyzed at day 61 post-immunization (Ctrl n = 3 mice, KO n = 3 mice). **(g)** GC B cell frequency. **(h)** B220^+^CD38^+^GL7^-^IgD^-^ MBC frequency. **(i)** CD138^+^B220^+^ plasmablast frequency. **(j)** Splenic CD138^+^B220^-^ PC frequency. **(k)** Bone marrow (BM) CD138^+^B220^-^ PC frequency. (two-sided unpaired *t*-test, bar plots show mean ± standard deviation)

MBC progenitors, pre-MBCs, differentiate from LZ B cells in the GC and express some MBC markers such as CCR6^48–50^. We observed a decreased frequency of pre-MBCs in the LZ of *Brwd1*-KO^GC^ mice (Extended Data Fig 5a-b). However, due to the increase in the total number of GC B cells in these mice, the number of pre-MBCs was comparable to control mice (Extended Data Fig 5c). A similar trend in pre-MBCs was observed in *Brwd1*-Het^GC^ mice (Extended Data Fig 5d-e). The MBC population is heterogeneous, and MBCs can differentiate from both activated B cells and GC B cells^51^. To measure GC-dependent MBCs, we examined multiple markers enriched in GC-derived MBCs, namely CD73, CD80, PD-L2, IgG1, and the absence of IgD^34,52–56^. Flow cytometry for various combinations of these markers showed no difference in the GC-derived MBC subsets (Extended Data Fig 5f-k).

To determine whether *Brwd1*-KO^GC^ mice produce long-lived PCs in the BM and mount a memory recall response, we immunized mice with SRBCs conjugated to 4-hydroxy-3-nitrophenylacetyl (NP) followed by a boost immunization at day 5. On day 56, we immunized again with NP-SRBCs, and on day 61 we measured B cell populations in the spleen and bone marrow (Fig 5f). There were not significant differences in the number or frequency of CD38^+^GL7^-^IgD^-^ MBCs, GL7^+^ GC B cells, CD138^+^B220^+^ plasmablasts, or CD138^+^B220^-^ PCs (Fig 5g-j). Similarly, we observed no significant difference in bone marrow CD138^+^ PCs (Fig 5k). In total, these results indicate that BRWD1 is not important for differentiation of GC-dependent MBCs, PCs, or plasmablasts.

### BRWD1 is necessary for optimal affinity maturation

We next explored whether BRWD1 is important for the principal functions of the GC, SHM and affinity maturation. We immunized mice with NP bound to keyhole limpet hemocyanin (KLH), which elicits a B cell response in which most antibodies specific to NP are of the IgG1 isotype and include the V_H_186.2 variable heavy chain gene segment^57^ (Fig 6a). We used *Aicda^Cre/wt^* ROSA-LSL-tdTomato^fl/fl^ mice to control for *Aicda* heterozygosity. At day 14 post-immunization, we sorted tdT^+^ GC B cells by FACS and cloned the V_H_186.2 segment for sequencing^57^. To prevent contamination by plasma cells, which have high levels of *Ig* transcripts, we used a primer specific to the Cγ1-membrane-encoding exon^57^.

**Figure 6.**
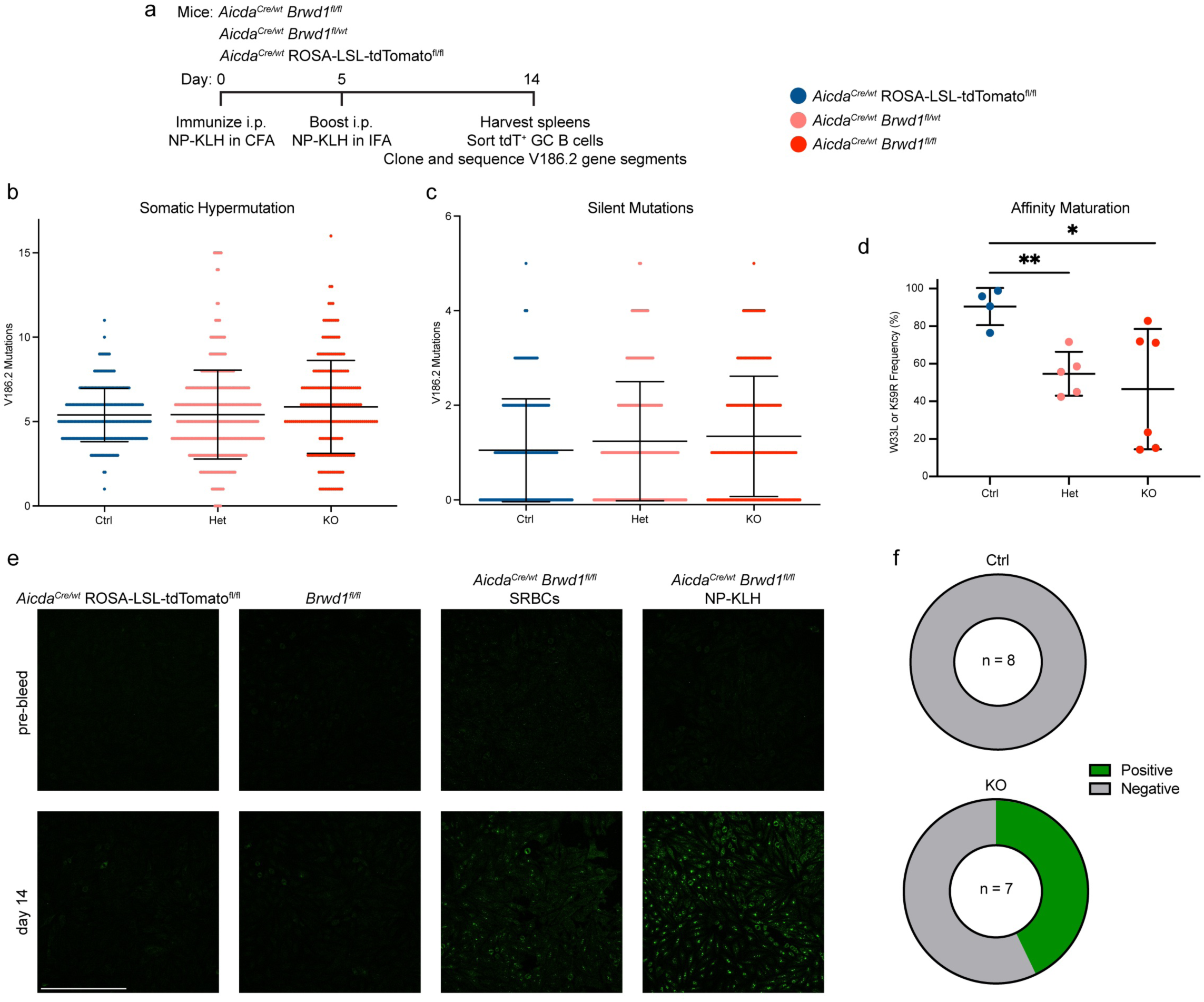
BRWD1 enhances affinity maturation. ***(a****) Aicda^Cre/wt^ Brwd1^fl/fl^* (KO), *Aicda^Cre/wt^ Brwd1^fl/wt^* (Het), and *Aicda^Cre/wt^* ROSA26-LSL-tdTomato^fl/fl^ (Ctrl) mice were immunized i.p. with NP-KLH in complete Freund’s adjuvant (CFA) and boosted with NP-KLH in incomplete Freund’s adjuvant (IFA) at day 5. At day 14 post-immunization, spleens were collected, tdT^+^ GC B cells were sorted, and V_H_186.2 gene segments were cloned and sequenced (Ctrl n = 4 mice, Het n = 5 mice, KO n = 6 mice). **(b)** Frequency of mutations within V_H_186.2 in GC B cells. **(c)** Frequency of silent mutations within V_H_186.2 in GC B cells. **(d)** Frequency of W33L or K59R amino acid substitutions that increase the affinity of V_H_186.2 to NP. **(e-f)** Control and knockout mice were immunized with NP-KLH in CFA or with SRBCs (KO n = 7 mice, Ctrl n = 8 mice). Sera was collected before immunization or at day 14 post-immunization and used to stain HEp-2 cells. (e) Representative images from a control *Aicda^Cre/wt^* ROSA-LSL-tdTomato^fl/fl^ mouse, a control *Brwd1^fl/fl^* mouse, a knockout *Aicda^Cre/wt^ Brwd1^fl/fl^* mouse immunized with SRBCs, and a knockout *Aicda^Cre/wt^ Brwd1^fl/fl^* mouse immunized with NP-KLH with CFA. Scale bar = 300 µm. (f) Proportion of mice with positive anti-nuclear antibody tests. Control *Brwd1^fl/fl^* and *Aicda^Cre/wt^* ROSA-LSL-tdTomato^fl/fl^ mice are grouped. Mice immunized with either SRBCs or NP-KLH are grouped. (\**p* < 0.05, \*\**p* < 0.01, two-sided unpaired *t*-test, lines show mean ± standard deviation)

Sequencing of V_H_186.2 revealed that tdT^+^ GC B cells from *Brwd1*-KO^GC^ mice and *Brwd1*-Het^GC^ mice undergo the same rate of SHM as GC B cells from *Aicda^Cre/wt^* ROSA-LSL-tdTomato^fl/fl^ control mice (Fig 6b). The rate of silent SHM, which is not affected by selection, was also similar between KO, Het, and control GC B cells (Fig 6c). In contrast to control GC B cells, KO and Het GC B cells had a significantly broader distribution of SHM with some clones containing a high number of mutations and some clones with none or one mutation (F test, *p* < 0.0001). In clones with a high amount of SHM, mutations were distributed throughout V_H_186.2 (Extended Data Fig 6a). Furthermore, the increased SHM distribution relative to controls was consistent across individual mice (Extended Data Fig 6b). The rate of mutations within framework regions and complementarity determining regions was the same between groups (Extended Data Fig 6c-e). Furthermore, there was no significant difference between groups in the rate of unproductive immunoglobulin mutations (Extended Data Fig 6f). In the WT GC cycle, there are relatively fixed, deterministic relationships between selection, proliferation, and SHM^20,23,27,58,59^. Our results suggest that upon loss of *Brwd1*, these tight relationships are uncoupled and the degree of proliferation following SHM and selection becomes stochastic.

We also measured affinity maturation by comparing the frequency of mutations known to increase affinity toward NP including W33L and K59R^60^. KO and Het GC B cells had an approximately two-fold decrease in the frequency of high affinity mutations relative to control GC B cells (Fig 6d). These results demonstrate that BRWD1 is necessary for optimal affinity maturation of GC B cells. Furthermore, decreased high affinity mutations in the Het GC B cells shows that the amount of BRWD1 is important for affinity maturation.

In addition to selection for higher affinity, GCs also eliminate autoreactive clones that arise from SHM^61–64^. To test GC negative selection, we collected sera from mice immunized with SRBCs or NP-KLH and tested reactivity to HEp-2 cells (Fig 6e). Prior to immunization, sera from all mice did not bind HEp-2 cells. Likewise, there was no detectable HEp-2 reactivity in control mouse serum after immunization. In contrast, sera from *Brwd1*-KO^GC^ mice immunized with either SRBCs or NP-KLH bound to HEp-2 cell nuclei, indicating self-reactivity. Approximately 40% of *Brwd1*-KO^GC^ mice demonstrated HEp-2 reactivity (Fig 6f). These results suggest that *Brwd1*-KO^GC^ mice are beginning to break tolerance following immunization, although the self-reactive sera may not be pathologically significant.

### BRWD1 is required for GC B cell subset transcriptional identity

We next performed RNA-seq on GC B cells from the LZ, DZp, and DZd of *Brwd1*-KO^GC^ mice and compared them with WT GC B cell subsets. PCA of these populations revealed that *Brwd1*^-/-^ GC B cells were transcriptionally distinct from WT GC B cells (Fig 7a-b). Comparison of WT and KO GC B cells within the same subset revealed a large transcriptional increase within the *Brwd1*^-/-^ LZ, DZp, and DZd (Fig 7c-e). Interestingly, many of the same genes were upregulated across subsets (Fig 7f, Extended Data Fig 7a-f). For example, 57% of genes increased in *Brwd1*^-/-^ LZ cells were also increased in *Brwd1*^-/-^ DZp and DZd cells (Extended Data Fig 7b). A heatmap of differential expression between all groups revealed many genes with similar expression in each *Brwd1*^-/-^ GC B cell subset (Extended Data Fig 7g). This was especially true of *Brwd1*^-/-^ DZp and DZd cells. Together, these results demonstrate that BRWD1 represses a large transcriptional program that is shared across *Brwd1*^-/-^ GC B cell subsets.

**Figure 7.**
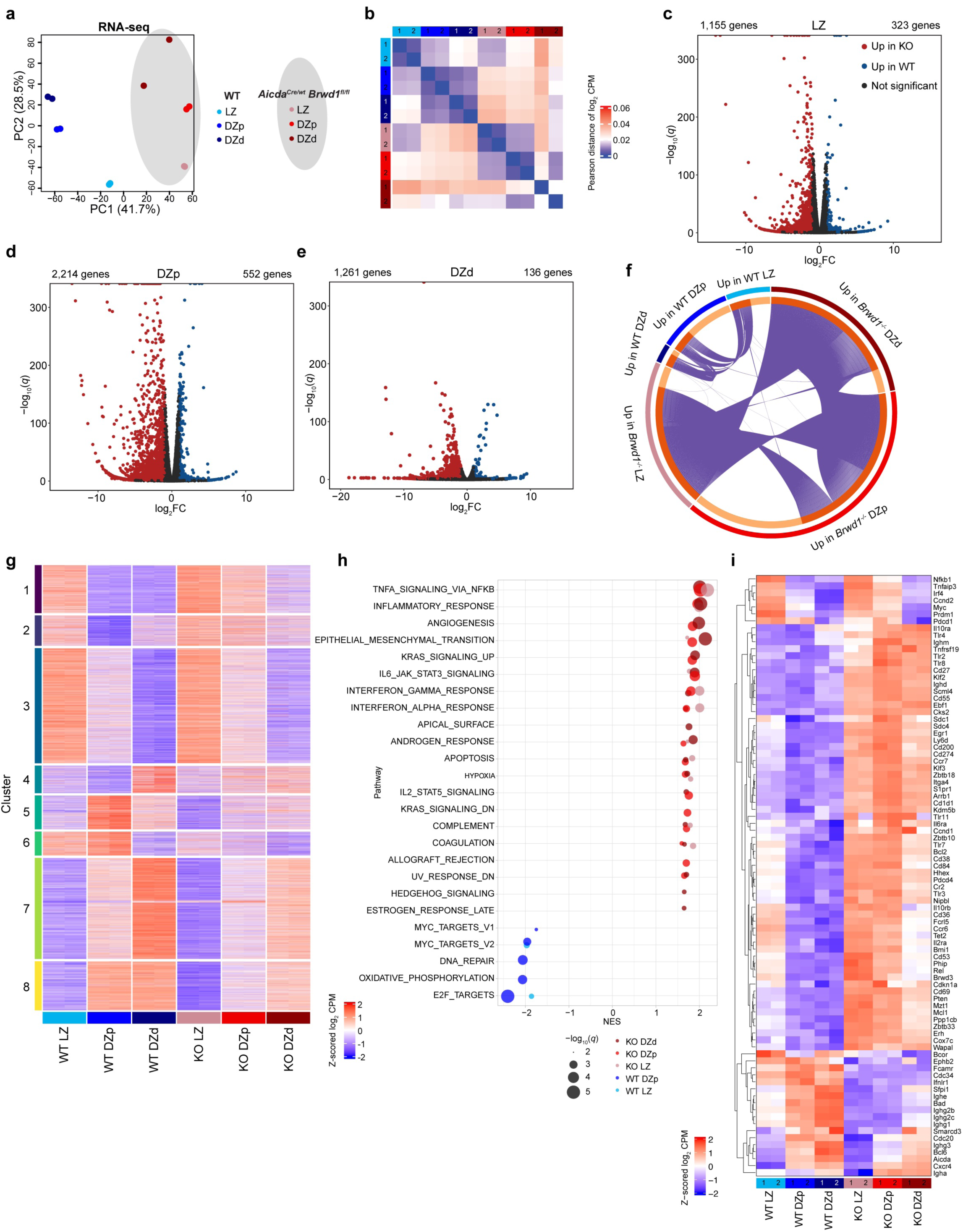
BRWD1 is required for GC B cell subset transcriptional identity. **(a)** PCA plot of RNA-seq of each GC subset in WT and *Aicda^Cre/wt^ Brwd1^fl/fl^* (KO) mice. n = 2 per cell type. Each n represents cells pooled from 20 mice. WT GC B cell RNA-seq was previously published^14^. **(b)** Heatmap of Pearson correlation of RNA-seq. **(c-e)** Volcano plots of differentially expressed genes (log_2_ fold change >1, *q* < 0.05) between KO and WT GC B cells from the LZ (c), DZp (d), and DZd (e). **(f)** Differentially expressed genes between *Brwd1^-/-^* and WT GC B cells from the LZ, DZp, or DZd. Purple lines show genes shared between groups. Dark orange signifies genes in the group that are shared with another group. **(g)** Gene expression of WT and KO GC subsets at clusters generated by unsupervised *K*-means clustering of WT GC subsets, performed previously^14^. Z-scored log_2_ counts per million (CPM) are plotted. **(h)** Gene set enrichment analysis (GSEA) using the Molecular Signatures Database hallmark genes sets for each GC subpopulation. Pathways are ordered by normalized enrichment score (NES). Pathways with *q* < 0.01 are shown. **(i)** Z-scored log_2_ CPM of notable genes expressed across the LZ, DZp, and DZd of WT and KO mice.

In WT GC subsets, RNA expression clustered into 8 groups, which previously revealed a transcriptional program unique to each GC subset, most notably the DZp (Fig 7g)^14^. Surprisingly, *Brwd1*^-/-^ DZp cells lost this unique transcriptional program and instead had intermediate expression between *Brwd1*^-/-^ LZ and DZd cells. For example, WT DZp cells had the greatest expression of genes in clusters 5 and 6 and the lowest expression of genes in clusters 1 and 2 (Extended Data Fig 7h). Yet *Brwd1*^-/-^ DZp cells had gene expression intermediate that of *Brwd1*^-/-^ LZ and DZd cells for genes in these same clusters. Thus, BRWD1 specifically maintains the DZp transcriptional program.

To understand the function of dysregulated genes in *Brwd1*^-/-^ GC B cells, we performed a gene set enrichment analysis (GSEA) comparing WT and *Brwd1*^-/-^ GC B cells from each GC subset. GSEA revealed a significant enrichment of many pathways in *Brwd1*^-/-^ GC B cells including TNFα signaling via NFκB and the inflammatory response (Fig 7h, Extended Data Fig 7i-n). Similarly, a gene ontology (GO) pathway analysis of genes significantly upregulated in WT or *Brwd1*^-/-^ GC B cells from each GC subset also revealed shared transcriptional pathways in *Brwd1*^-/-^ GC B cells, including multiple immune and inflammatory response GO pathways (Extended Data Fig 7o). Within these pathways were many differentially expressed genes important for GC B cell biology (Fig 7i). Taken together, these results demonstrate that within the genes upregulated in each *Brwd1*^-/-^ GC B cell subset are inflammatory transcriptional programs not expressed in WT GC B cells.

RNA-seq revealed increased expression of the *Brwd1* homologs *Phip* (also known as *Brwd2*) and *Brwd3* in the *Brwd1*^-/-^ GC subsets (Extended Data Fig 7p). These genes could partially compensate for *Brwd1* deletion in *Brwd1*-KO^GC^ mice, thereby mitigating the observed phenotype.

### BRWD1 maintains chromatin accessibility differences across GC subsets

Finally, we performed ATAC-seq on LZ, DZp, and DZd GC B cells from *Brwd1*-KO^GC^ and WT mice to understand the effects of deleting *Brwd1* in GC B cells. Comparison with WT GC B cells revealed that *Brwd1*^-/-^ GC B cells were epigenetically distinct in the PCA space (Fig 8a). In *Brwd1*^-/-^ GC B cells, differences in chromatin accessibility were diminished as shown by PCA where *Brwd1*^-/-^ samples clustered more closely together. Indeed, the biological coefficient of variation (BCV) was lower across *Brwd1*^-/-^ GC subsets (12.7%) compared to WT GC subsets (25.2%). In WT GC B cells, many accessibility peaks significantly changed during transitions between subsets; however, these changes were diminished in *Brwd1*^-/-^ GC B cells (Fig 8b). For example, transition between the DZd and LZ involves 61,451 differential accessibility peaks in WT cells but only 15,522 differential accessibility peaks in *Brwd1*^-/-^ GC B cells. A striking example of this occurred at the *Myc* locus. In WT cells, accessibility peaks at *Myc* and downstream enhancers in the LZ were closed in DZd GC B cells. In contrast in *Brwd1*^-/-^ GC B cells, chromatin accessibility was unchanged at the *Myc* locus across GC B cell subsets (Fig 8c).

**Figure 8.**
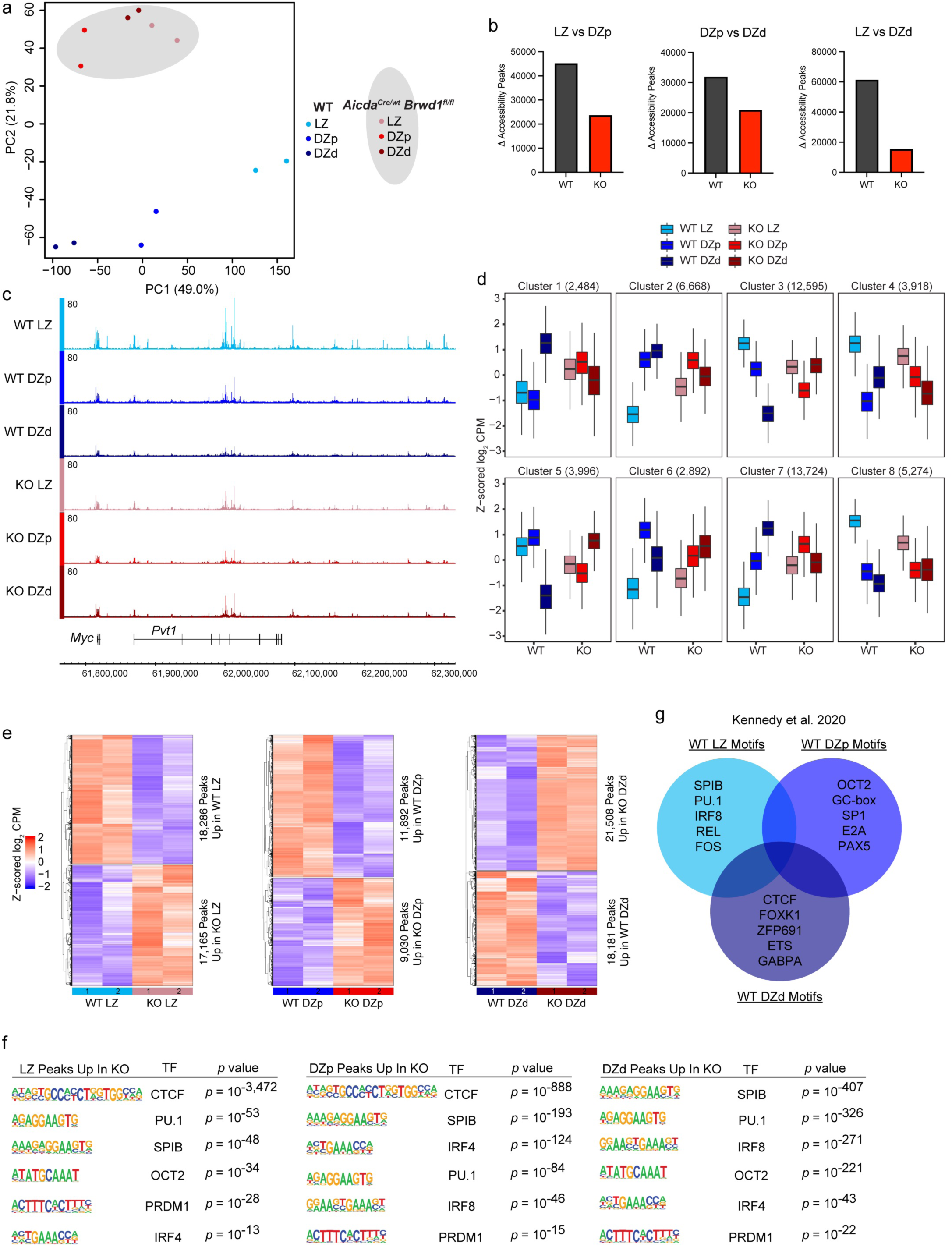
BRWD1 maintains the distinct epigenetic states of each GC subset. **(a)** PCA plot of ATAC-seq of each GC subset in WT and *Aicda^Cre/wt^ Brwd1^fl/fl^* (KO) mice. n = 2 per cell type. Each n represents cells pooled from 20 mice. WT GC B cell ATAC-seq was previously published^14^. **(b)** Number of differentially regulated accessibility peaks (*q* < 0.05) between GC subsets in WT and KO mice. **(c)** Chromatin accessibility tracks at the *Myc* locus (mm9, chromosome 15) for each GC subset in WT and KO. **(d)** Chromatin accessibility of WT and KO GC subsets at clusters generated by unsupervised *K*-means clustering of WT GC subsets, performed previously^14^. Z-scored log_2_ counts per million (CPM) are plotted. Box limits show interquartile range (IQR), and center lines show median. Maximum and minimum values (whiskers) are defined as Q3 + 1.5 x IQR and Q1 – 1.5 x IQR, respectively. **(e)** Heatmaps of differentially accessible peaks (*q* < 0.05) between KO and WT GC B cells from the LZ, DZp, or DZd. Z-scored log_2_ CPM are plotted. **(f)** TF motifs enriched in accessible regions that are up in KO GC B cell subsets (*q* < 0.05) generated using HOMER. **(g)** Venn diagram of enriched TF motifs in each WT GC subset previously described^14^.

In WT GC B cells, accessibility peaks clustered into eight groups each with a different distribution pattern across GC subsets (Fig 8d)^14^. In contrast, *Brwd1*^-/-^ GC subsets had decreased variability within these groups compared to WT. Accessibility was especially dysregulated in *Brwd1*^-/-^ DZp and DZd cells. For example, at sites in cluster 5, WT DZd cells have low accessibility; however, *Brwd1*^-/-^ DZd cells had greater accessibility than *Brwd1*^-/-^ LZ or DZp cells at these same sites. In a similar manner for cluster 6, WT DZp cells have the greatest accessibility; however, these sites were accessible in both *Brwd1*^-/-^ DZp and DZd cells. Generally, sites uniquely accessible in WT LZ cells were also accessible in *Brwd1*^-/-^ LZ cells (clusters 3, 4, and 8), yet these clusters also showed a blending across *Brwd1*^-/-^ subsets. These results demonstrate that BRWD1 is necessary for polarizing the chromatin accessibility states of GC B cells.

Comparison of WT and *Brwd1*^-/-^ cells within the same GC B cell subset revealed large differences in chromatin accessibility (Fig 8e). Analysis of the chromatin accessibility peaks that were increased in *Brwd1*^-/-^ cells in each subset revealed similar TF motifs across subsets (Fig 8f). We previously reported that each GC subset is associated with unique TF motif accessibility (Fig 8g)^14^; however, *Brwd1*^-/-^ GC B cell subsets were enriched for many of the same TF motifs. For example, in WT cells CTCF is most enriched in DZd cells^14^, yet in *Brwd1*^-/-^ GCs the CTCF motif was enriched in LZ and DZp cells. Similarly, OCT2 is the top enriched motif in WT DZp cells^14^, yet this motif was enriched in *Brwd1*^-/-^ cells in the LZ and DZd. Across all three KO GC subsets, the accessibility at motifs for PU.1, SPIB, PRDM1, and IRF4 was shared while in WT cells, accessibility at each site was restricted to one GC subset. In total, we conclude that BRWD1 is necessary for maintaining the unique transcriptional and epigenetic states of each GC subset.

## DISCUSSION

There are four stages of B cell development and activation in which the molecular necessities of diversity must be insulated from, and coordinated with, proliferation and selection^15^. The precise cellular states to which light chain recombination and SHM are restricted have been defined^14,30,31^. In contrast, we only know the general developmental windows in which heavy chain recombination and CSR occur. BRWD1, which is first highly expressed in small pre-B cells, orchestrates the large epigenetic and transcriptional transitions that enable and insulate light chain recombination, CSR, and SHM from proliferation^30,31^. Therefore, BRWD1 is essential for most of the defining molecular events of B cell adaptive immunity.

Deletion of BRWD1 in Fol B cells suggests that it facilitates molecular transitions rather than maintaining cell states. Indeed, BRWD1 did not substantially contribute to the transcriptional state of Fol B cells. Rather, it primed the epigenome by both opening binding sites for TFs necessary for GCs while repressing Ets1 binding sites, a TF that can inhibit B cell differentiation^41–44^. These results are consistent with prior work in B cell development where BRWD1 mediates transitions between cell states by priming cells for the next stage of differentiation^30,31^. Indeed, our data supports a general model that throughout the B cell lineage the opening and closing of TF motifs precede cellular differentiation^65^.

In addition to preparing the epigenetic landscape for GC differentiation, BRWD1 enabled CSR by remodeling the 3D organization of the class switch locus. BRWD1 binds to topologically associating domain (TAD) boundaries where it converts static to dynamic cohesin to initiate intra-TAD loop extrusion^32^. The class switch locus had the hallmarks of cohesin conversion, including accumulation of static cohesin (RAD21 and SMC3) at TAD boundaries and loss of dynamic cohesin (co-occurring RAD21, SMC3, NIPBL, and WAPL) in *Brwd1^-/-^* cells. However, BRWD1 appeared to convert cohesin at the opposite TAD boundary to which it was bound. This suggests that BRWD1 can convert distant cohesin on the same chromosome if this static cohesin is within genomic distances accessible by diffusion^36^.

In GC B cells, BRWD1 appeared to maintain GC epigenetic identity with a striking collapse of unique TF binding sites toward a common state among *Brwd1*^-/-^ GC B cell subsets. Given the functions of BRWD1 in other contexts, we postulate that this epigenetic blending reflected a failure to prime transitions between GC B cell subset states. This epigenetic blending did not precisely map to transcriptional changes. There was blending of transcriptional programs, especially those associated with *Myc* and proliferative programs, across *Brwd1*^-/-^ GC subsets. However, prominent inflammatory gene signatures were also induced in all *Brwd1*^-/-^ GC subsets. The exact mechanism of this latter induction is not clear. We did not consistently observe an upregulation of apoptotic pathways, and we failed to detect an increase in apoptotic cells (data not shown). Regardless of mechanism, inflammatory signals could subvert both GC positive and negative selection by providing activation signals not determined by antigen avidity.

Indeed, we observed specific defects in both affinity maturation and tolerance. The defect in affinity maturation could be due to dysregulation of either positive or negative selection of *Brwd1^-/-^* GC B cells. For example, *Brwd1^-/-^* GC B cells with a low affinity for NP may still proliferate and cycle through the GC with diminished or no positive selection signals. Alternatively, it is possible that the negative selection signals to either exit the GC or undergo apoptosis are not properly received in these cells.

While *Brwd1*^-/-^ GCs had normal SHM rates, the distribution of SHM in GC B cells was greater such that some clones had few if any mutations while others had 10 or more mutations at 14 days post-immunization. Normally, there is a stringent relationship between SHM, selection, and proliferation with B cells proliferating once or twice following LZ selection^20,23,27,58,59^. In contrast, our data are consistent with *Brwd1*^-/-^ GC B cells proliferating chaotically following selection.

Surprisingly, BRWD1 was only necessary for GC selection. In fact, many characteristics of the GC response, including zonal organization, MBC and PC differentiation, and Tfh cells, were normal in *Brwd1*-KO^GC^ mice. These observations are consistent with what is known about GC evolutionary biology. GCs and affinity maturation convergently evolved only in endothermic mammals and birds approximately 200 million years ago^66^. In contrast, SHM arose approximately 450 million years ago in ectothermic cartilaginous fish, which can also generate MBCs and PCs^66–68^. In cartilaginous fish, SHM functions to diversify antibody responses and only modestly enhances affinity^69,70^. Efficient affinity maturation only developed with the advent of GCs. It has been argued that GCs evolved to deal with the more rapid cell proliferation of endotherms^66^. Indeed, we postulate that with higher proliferative rates, insulation between SHM and mitosis via specialization of GC subsets became an evolutionary imperative to diminish the risk of neoplastic transformation. These observations suggest that BRWD1 selectively mediates functions that were only acquired with the evolutionary advent of GCs.

We have demonstrated that BRWD1 mediates important B cell fate transitions. In both B cell development and the GC, BRWD1 represses *Myc*^31^. However, the genetic programs regulated by BRWD1 at each stage are not identical. An obvious example is that BRWD1 mediates *Igk* contraction in small pre-B cells yet contributes to CSR in the periphery. BRWD1 is recruited to specific histone modifications downstream of signaling pathways^29,30^. Therefore, it is likely that signaling and developmental contexts dictate where BRWD1 is recruited in the genome and which TADs and genes it acts on.

Considering BRWD1’s essential role in chromatin loop extrusion, it was surprising that *Brwd1*-KO^Fol^ and *Brwd1*-KO^GC^ mice still formed GCs. One possibility is that *Brwd1* homologs *Phip* and *Brwd3* compensated for *Brwd1* loss, although these homologs may lack functional domains unique to *Brwd1* (data not shown). Alternatively, our results may reveal the degree to which loop extrusion is necessary for cell differentiation. Disruption of loop extrusion by depleting cohesin in embryonic stem cells only modestly effects transcription^71,72^. However, loop extrusion by cohesin is required for many promoter-enhancer interactions, especially those occurring over longer distances in differentiated cells^73,74^. Our results suggest that BRWD1-mediated changes in chromatin topology allow GC B cell subsets to fully polarize as they rapidly cycle through the GC.

In conclusion, our results show how BRWD1 is necessary for GC integrity. Deletion of *Brwd1* in GC B cells resulted in major transcriptional and epigenetic disruption, which had specific effects on GC function, namely proliferation, affinity maturation, and tolerance. While we previously proposed a three-zone model of the GC that illuminates the transcriptional and epigenetic variation throughout the GC cycle^14^, our data here validate this model and demonstrate that the molecular distinctions between the three GC zones are essential for GC function.

## METHODS

### Mice

*Brwd1*-floxed mice were generated by Ingenious Targeting Laboratory. A targeting vector was designed containing a Lox71 site 5’ of *Brwd1* exon 6, a neomycin-resistance cassette flanked by flippase recognition target (FRT) sites between *Brwd1* exons 7 and 8, an inverse tdTomato reporter with a bovine growth hormone polyadenylation sequence (BGHpA), and a Lox66 site 3’ of *Brwd1* exon 8. The Lox72 and Lox66 sites were in opposite orientation to one another. The targeting vector was introduced by electroporation to C57BL/6 embryonic stem cells containing flippase. Resulting cells were microinjected into Balb/c blastocysts. Resulting chimeras with a high percentage black coat color were mated to C57BL/6 WT mice to generate germline neomycin-resistance cassette-deleted mice. PCR and sequencing were used to confirm the deletion of the neomycin-resistance cassette, the presence of the tdTomato cassette, and the presence of the Lox66 site. PCR was used to confirm the absence of the flippase transgene and the presence of the Lox71 site.

Routine genotyping of the *Brwd1-floxed* mice was done with the LOX1 and SDL2 primers, which cover the 5’ Lox71 site (Supplementary Table 1). PCR conditions were 94 °C for 2 min, then 35 cycles of 94 °C for 30 s, 60 °C for 30 s, and 72 °C for 1 min, followed by 72 °C for 2 min. The WT amplicon is 389 bp and the floxed amplicon is 431 bp.

Wild type C57BL/6J mice (stock #000664), *Aicda^Cre^* mice (stock #007770), and Ai14 Rosa-CAG-LSL-tdTomato-WPRE mice (stock #007914) were purchased from Jackson Laboratory. *Cd23^Cre^* mice were obtained from the laboratory of Jayanta Chaudhuri (Memorial Sloan Kettering). Mice were housed in the University of Chicago animal facilities, and studies were performed in accordance with the guidelines of the Institutional Animal Care and Use Committee (protocol number 71577). Female and male mice were used at 6-12 weeks of age.

### Immunizations

Mice were immunized intraperitoneally with 10^9^ sheep red blood cells (SRBCs, Lampire Biological Laboratories) in PBS and boosted with 10^9^ SRBCs 5 days later. Alternatively, mice were immunized intraperitoneally with 200 µl of 4-hyrdoxy-3-nitrophenyl-acetyl (NP)-keyhole limpet hemocyanin (KLH) (1 mg/ml, valency of 27 or 33) in a 1:1 ratio with Complete Freund’s Adjuvant (CFA)^57^. Mice were boosted with NP-KLH in a 1:5 ratio with Incomplete Freund’s Adjuvant (IFA) at day 5. Mice were also immunized intraperitoneally with 10^9^ SRBCs conjugated with NP by incubating NP-OSu in 0.15 M NaHCO_3_ with SRBCs. For this experiment, mice were boosted with NP-SRBCs at days 5 and 56 post-immunization.

### Flow cytometry

On day 5 or 14 post-immunization, spleens were harvested, and cells were resuspended in 1X phosphate-buffered saline (PBS) with 3% (v/v) fetal bovine serum (FBS). Erythrocytes were lysed with ACK lysis buffer (Lonza). Splenocytes were passed through a cell strainer to obtain single cell suspensions. Fc block was done with anti-mouse-CD16/CD32 (2.4G2). Cells were stained with viability dye eFluor506 (eBioscience).

Fol B cells were stained with anti-B220-APC/Cy7 (RA3-6B2), anti-CD19-PerCP/Cy5.5 (1D3), anti-CD93-BV421 (AA4.1), anti-CD23-PE/Cy7 (B3B4), and anti-CD21-APC (7G6). GC B cells were stained with anti-B220-AF488 (RA3-6B2), GL7-PerCP/Cy5.5, anti-CD95-BV421 (Jo2), anti-CD83-PE/Cy7 (Michel-19), and anti-CXCR4-APC (2B11). Cells were also stained with various combinations of anti-CD138-APC (281-2), anti-IgD-PE/Cy7 (11-26c), anti-BCL6-PE/Cy7 (K112-91), Annexin V-FITC (BD Biosciences), FxCycle Violet (Invitrogen), and the FAM FLICA poly caspase kit (Bio-Rad). Cells were fixed with the Foxp3/Transcription factor staining buffer kit (eBioscience). Tfh cells were stained with anti-CXCR5-BV421 (L138D7), anti-PD-1-PerCP/Cy5.5 (RMP1-30), anti-CD4-APC (RM4-5), and anti-CD3-FITC (145-2C11).

For flow cytometry of MBC subsets, cells were stained with anti-CD38-BUV496 (90/CD38), anti-IgM-BUV615 (II/41), anti-PD-L2-BUV737 (TY25), anti-CCR6-BV421 (29-2L17), anti-IgD-BV480 (11-26c.2a), anti-CD80-BV650 (16-10A1), anti-IgG1-BV711 (A85-1), anti-CD95-BV786 (Jo2), anti-B220-AF488 (RA3-6B2), GL7-PerCP/Cy5.5, anti-CD83-PE/Cy7 (Michel-19), anti-CXCR4-APC (2B11), anti-CD138-APC-R700 (281-2), anti-CD73-APC-Fire750 (TY/11.8), and viability dye eFluor506 (eBioscience) with Brilliant Stain Buffer Plus.

Flow cytometry was done with a LSRFortessa (BD Biosciences) running FACSDiva version 8.0.2 and spectral flow cytometry was done with a Cytek Aurora. Flow cytometry data was analyzed with FlowJo version 10.8.1 (BD Biosciences).

### Cell sorting

Fol B cells were sorted by staining with anti-B220-BV421 (RA3-6B2), anti-CD93-FITC (AA4.1), anti-CD21-APC (7G6), and anti-CD23-PE/Cy7 (B3B4). Enrichment of GC B cells for fluorescence-activated cell sorting (FACS) was done using magnetic-activated cell sorting (MACS), as previously described^75^. GC B cell enrichment was done with anti-CD43-biotin (S7), anti-CD11c-biotin (N418), anti-IgD-biotin (11-26c.2a), streptavidin microbeads (Miltenyi Biotec), and LS columns (Miltenyi Biotec). FACS was performed on a FACSAria Fusion (BD Biosciences) running FACSDiva version 8.0.2.

### Microscopy

On day 14 post-immunization, spleens were harvested, transferred to OCT (Fisher HealthCare) and frozen on dry ice in 2-methylbutane. Frozen spleens were sectioned at 6 µm. For immunofluorescence microscopy, sections were fixed with 4% paraformaldehyde, permeabilized with 1X NP40 Permeating Solution in PBS (Boston Products), and Fc receptors were blocked with anti-mouse-CD16/CD32 (2.4G2, 1:200) in 10% normal donkey serum (Jackson ImmunResearch).

For *Cd23^Cre/wt^ Brwd1^fl/fl^* mice, spleens were collected day 14 post-immunization. Tissue was stained with GL7-AF647 (1:20) in 5% normal donkey serum. Images were acquired on a Leica Stellaris 8 laser scanning confocal microscope (Leica Microsystems) at 10X magnification.

For *Aicda^Cre/wt^ Brwd1^fl/fl^* mice, tissue was stained with GL7-AF647 (1:20), anti-CD4-AF594 (GK1.5, 1:20), anti-CD35-BV421 (8C12, 1:20), anti-IgD-BV711 (11-26c.2a, 1:20), anti-tdTomato (polyclonal, Rockland, 1:50), followed by donkey anti-rabbit-IgG-AF488 (polyclonal, Invitrogen, 1:1000) in 5% normal donkey serum. The GL7 channel and differential interference contrast (DIC) were collected at 10X magnification, and all channels were collected for individual GCs at 20X magnification.

### Image analysis

Images were visualized and analyzed with Fiji version 2.9.0^76^. For quantification of GC size, GL7 images were Gaussian filtered (sigma = 10), and an intensity threshold and size exclusion threshold were used to remove background signal.

For mean pixel intensity (MPI) image analysis, images collected at 20X magnification were normalized to the 99^th^ percentile for each channel. For a given channel, each image in the dataset was standardized through histogram matching to ensure similar contrast throughout the dataset. GCs were segmented by applying a Gaussian filter (sigma = 5) to the GL7 channel, then adjusting contrast and thresholding. The binary mask created by this segmentation algorithm was separated by instance using the regionprops function in the sci-kit image python library^77^. After identifying discrete objects (GCs) from the binary segmentation mask, small objects were rejected using a size filter, and GCs at the image boundaries were rejected to ensure each GC was only measured once. GC area was computed from the resulting segmentation masks. The LZ was segmented by applying a similar Gaussian filtering and thresholding algorithm to the CD35 channel. Resulting LZ segmentations were masked using the GC mask defined above to ensure that the segmented LZ was fully encompassed within the GC segmentation. MPI (pixel value sum/segmented area) was calculated for each marker within the regions defined by the GC and LZ masks.

### *In vitro* class switch recombination

Splenic cells were harvested as above. B cells were enriched by incubating cells with anti-CD43-biotin (S7) and magnetic enrichment with LS columns (Miltenyi Biotec). For class switching to IgG1, cells were cultured in RPMI 1640 with L-Glutamine media (Gibco) with 10% FBS, 0.1% β-mercapto-ethanol, and 1% penicillin-strep at 37 °C, 5% CO_2_. For class switching to IgG1, 25 µg/ml LPS (Sigma-Aldrich) and 10 ng/ml IL-4 (R&D Systems) were added to the media.

After 96 hr, class switching was measured by flow cytometry. Cells were stained with anti-CD19-APC/Cy7 (1D3), anti-IgD-PE/Cy7 (11-26c.2a), anti-IgM-APC (II/41), anti-IgG1-FITC (RMG1-1), and eFluor506 viability dye (eBioscience).

### Somatic hypermutation

RNA was extracted from sorted GC B cells using the RNeasy Mini Kit (Qiagen). cDNA was generated using the SuperScript III First-Strand Synthesis System for RT-PCR kit (Invitrogen) and the mCy1-cDNA primer. A semi-nested PCR was performed using the V186.2-leader and V186.2-nested primers with the Cy1-PCR primer as has been described^57^. PCR products were purified with the MinElute PCR Purification Kit (Qiagen). Cloning was performed with the TOPO TA Cloning & Bacterial Transformation kit (Invitrogen). DNA from colonies was isolated with the PureLink Quick Plasmid Miniprep Kit (Invitrogen). DNA was sequenced with the M13 Forward primer.

The vector sequence was trimmed with 4Peaks (Nucleobytes). Alignment and mutations were analyzed with IMGT/V-QUEST^78^. Duplicate sequences from the same mouse, sequences that did not align most closely to V186.2, and sequences with an uneven distribution of mutations were not used. Mutations in CDR3 introduced by junctional diversity were not counted to measure SHM.

### Anti-nuclear antibody test

Mouse sera was diluted 1:100 in PBS. Kallestad HEp-2 cell line slides (BioRad) were incubated with the sera. Slides were stained with donkey anti-mouse-IgG-AF647 (1:1000) and Hoechst 33342 (1:1000). Immunofluorescence microscopy was done with a Dragonfly 200 confocal microscope (Andor).

### RNA-seq

Total RNA was isolated using a RNeasy kit (Qiagen). Libraries were prepared by the University of Chicago Genomics Facility before sequencing on the NovaSeq-X-Plus (Illumina). Raw reads were aligned to reference genome mm9 in a splice-aware manner using STAR^79^. Gene expression was quantified using featureCounts^80^ against UCSC genes, with Ensembl IG genes from mm10 converted to mm9 coordinates with UCSC liftOver.

### ATAC-seq

Cells were washed with PBS, then lysed with ATAC lysis buffer (10 mM Tris-HCl, pH 7.4, 10 mM NaCl, 3 mM MgCl2, 0.1% IGEPAL CA-630). Resulting nuclei were then incubated with tagmentation enzyme (Illumina). Libraries from purified samples were made with the Nextera Indexing kit (Illumina).

Raw reads were aligned to reference genome mm9 using BWA MEM^81^. Apparent PCR duplicates were removed using Picard MarkDuplicates (https://github.com/broadinstitute/picard/releases/ tag/2.11.0)

For ATAC-seq, read alignments were first adjusted to account for TAC transposon binding: +4 bp for +strand alignments, −5 bp for -strand alignments. The open chromatin enrichment track was generated by first creating a bedGraph from the raw reads using bedtools genomcov^82^, then converted to bigWig using UCSC tool bedGraphToBigWig. Tracks were normalized by the sum of alignment lengths over 1 billion. Open chromatin peaks were called using Macs2^83^ with no model set and no background provided. Peaks with a score greater than 5 were retained. To quantitatively measure changes in epigenetic enrichment, we first identified empirical regulatory elements based on the peak calls obtained from each sample in the analysis. Peaks were merged into a uniform set of regulatory elements using bedtools merge^82^. Enrichment levels for each regulatory element were then quantified with featureCounts^80^.

### Differential expression of RNA-seq and ATAC–seq

Differential expression statistics (fold-change and *p* value) were computed using edgeR on raw expression counts obtained from quantification (either genes or peaks)^84^. Pairwise comparisons were computed using exactTest, and multigroup comparisons using the generalized linear modeling capability in edgeR. In all cases, *p* values were adjusted for multiple testing using the FDR correction of Benjamini and Hochberg.

### Next generation sequencing analysis

RNA-seq, ATAC-seq, and ChIP-seq data was visualized with the Integrated Genome Browser^85^. Heatmaps were generated with ComplexHeatmap and Circlize in R, and plots were made with ggplot2^86,87^. Metascape data portal was used for pathway analyses and for RNA-seq circle plots^88^. Hypergeometric optimization of motif enrichment (HOMER) was used to perform known transcription factor binding motif analyses using the FindMotifsGenome function to generate enrichment and *p* values^89^. Gene set enrichment analysis (GSEA) was performed as described using the Molecular Signatures Database hallmark genes sets^90,91^. Hi-C data was visualized using Juicebox^92^. Hi-C arc plots were generated with HiCcompare by normalizing Hi-C matrices using the hic_loess function and calculating differences using the hic_compare function^93^. Then the plotBedpe function from Sushi.R was used to plot differences with *p* < 0.05^94^.

### Statistical analysis

Statistical analyses were performed with GraphPad Prism. Bar graphs are displayed as the mean ± standard deviation. Significance as defined by *p* value or false discovery rate (*q* value) are defined in the figure legends. All measurements were taken from distinct samples as described in the figure legends.

### Data availability

The data that support the findings of this study are available from the corresponding authors upon reasonable request. RNA-seq and ATAC-seq data were deposited in the Gene Expression Omnibus (GEO) database with accession code GSE264164. WT GC B cell RNA-seq and ATAC-seq data from GEO accession code GSE133743 was analyzed in this study^14^. Small pre-B cell Hi-C and ChIP-seq from GEO accession code GSE221519 was also analyzed^32^.

## Acknowledgments

This work was supported by National Institutes of Health (NIH) grants T32GM007281 (to N.E.W. and M.R.C.), F30 AI174324 (to N.E.W), R01 AI143778 (to M.R.C.), and AI150860 (to M.R.C. and M.M.). M.M.-C. is supported in part by NCATS through grant UL1TR002003. Flow cytometry and FACS was performed at the Cytometry and Antibody Technology Facility at the University of Chicago (RRID: SCR_017760) and next generation sequencing was done by the University of Chicago Genomics Facility (RRID: SCR_019196), both of which receive financial support from the Cancer Center Support Grant P30CA014599. Imaging was performed at the University of Chicago Integrated Light Microscopy Core (RRID: SCR_019197). Sanger sequencing was performed with the University of Chicago DNA Sequencing and Genotyping Facility. We thank Dr. Sandeep Gurbuxani for his help with analysis of H&E images.

## Author contributions

N.E.W., M.M., and M.R.C. conceived and designed experiments. N.E.W. performed and analyzed most of the experiments. D.E.K., M.L.V., Y.M.Y., and N.E.W. performed flow cytometry. J.A. and N.E.W. performed immunofluorescence microscopy. M.D. and N.E.W. analyzed immunofluorescence microscopy images. M.A. and N.E.W. performed *in vitro* class switch recombination experiments. J.V. assisted with SHM experiments. D.E.K. and N.E.W. performed RNA-seq and ATAC-seq. M.M.-C., D.E.K., and N.E.W. analyzed next generation sequencing data. M.M. analyzed the HiC data. N.E.W. wrote the first draft of the manuscript. N.E.W. and M.R.C. edited the manuscript.

---

**Extended Data Figure 1.**
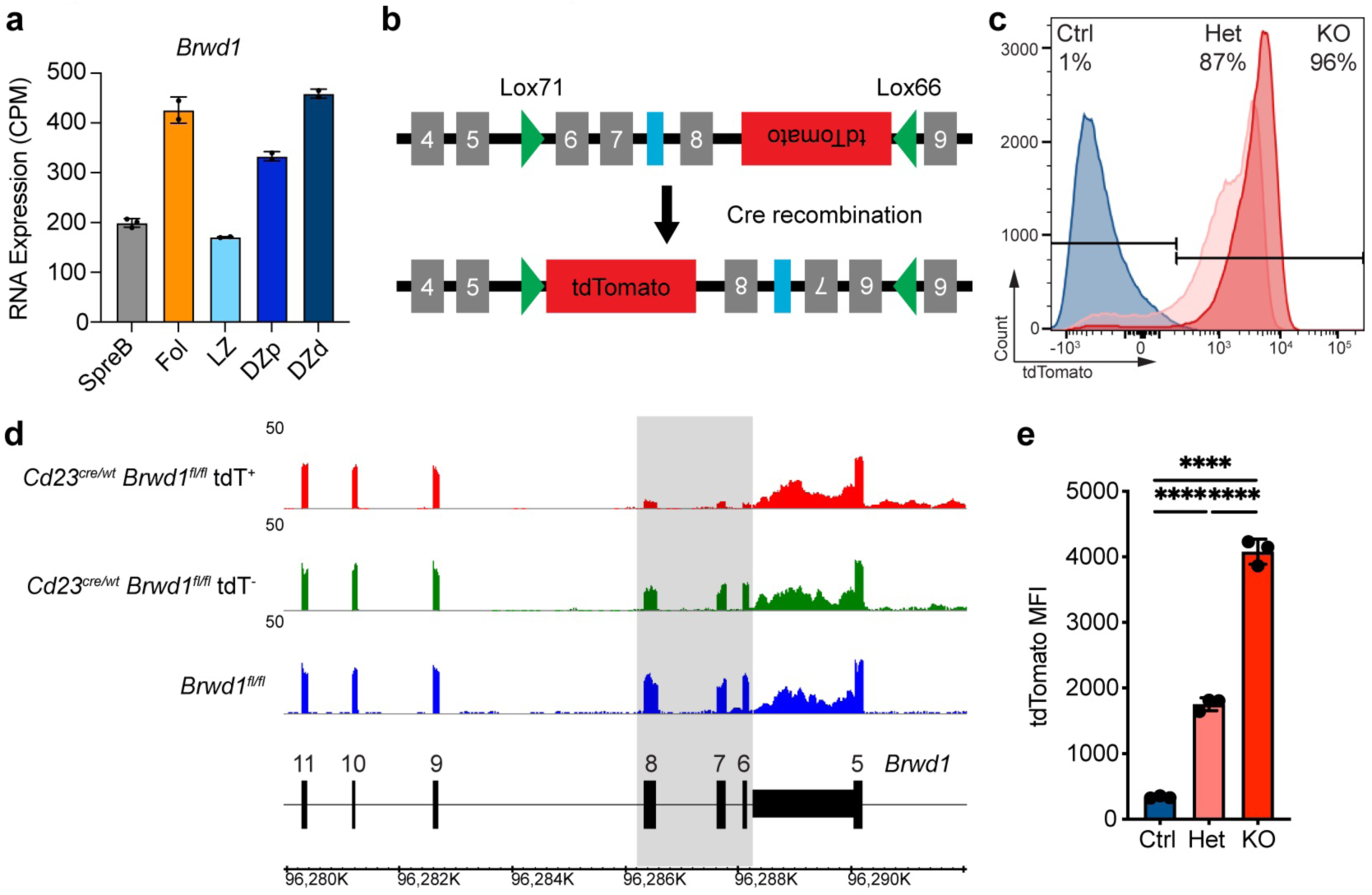
Design and characterization of *Brwd1*-floxed mouse. **(a)** RNA-seq of *Brwd1* expression for small pre-B, Fol B, and GC B cells in the LZ, DZp, and DZd (n = 3 for small pre=B cells, n = 2 for all other cell types). **(b)** Model of LoxP sites (green) surrounding exons 6, 7, and 8 of *Brwd1* with a tdTomato reporter in the inverse orientation. A single flippase recognition target (FRT) site (blue) remained following removal of a selection cassette. **(c)** tdTomato (tdT) expression by flow cytometry of splenic B220^+^CD19^+^CD93^-^ CD23^+^CD21^+^ Fol B cells in *Cd23^Cre/wt^ Brwd1^fl/fl^* (KO), *Cd23^Cre/wt^ Brwd1^fl/wt^* (Het), and *Brwd1^fl/fl^* (Ctrl) mice. Frequency of tdT^+^ Fol B cells is shown. **(d)** RNA-seq of *Brwd1* transcripts mapped to the *Brwd1* locus in sorted *Cd23^Cre/wt^ Brwd1^fl/fl^* tdT^+^, *Cd23^Cre/wt^ Brwd1^fl/fl^* tdT^-^, and *Brwd1^fl/fl^* Fol B cells. The region flipped in orientation following Cre recombination is highlighted grey. **(e)** Median fluorescent intensity (MFI) of tdT^+^ Fol B cells from *Cd23^Cre/wt^ Brwd1^fl/fl^* (KO), *Cd23^Cre/wt^ Brwd1^fl/wt^* (Het), and *Brwd1^fl/fl^* (Ctrl) mice (n = 3 mice per group). (\*\*\*\**p* < 0.0001, two-sided unpaired *t*-test, bar plots show mean ± standard deviation)

**Extended Data Figure 2.**
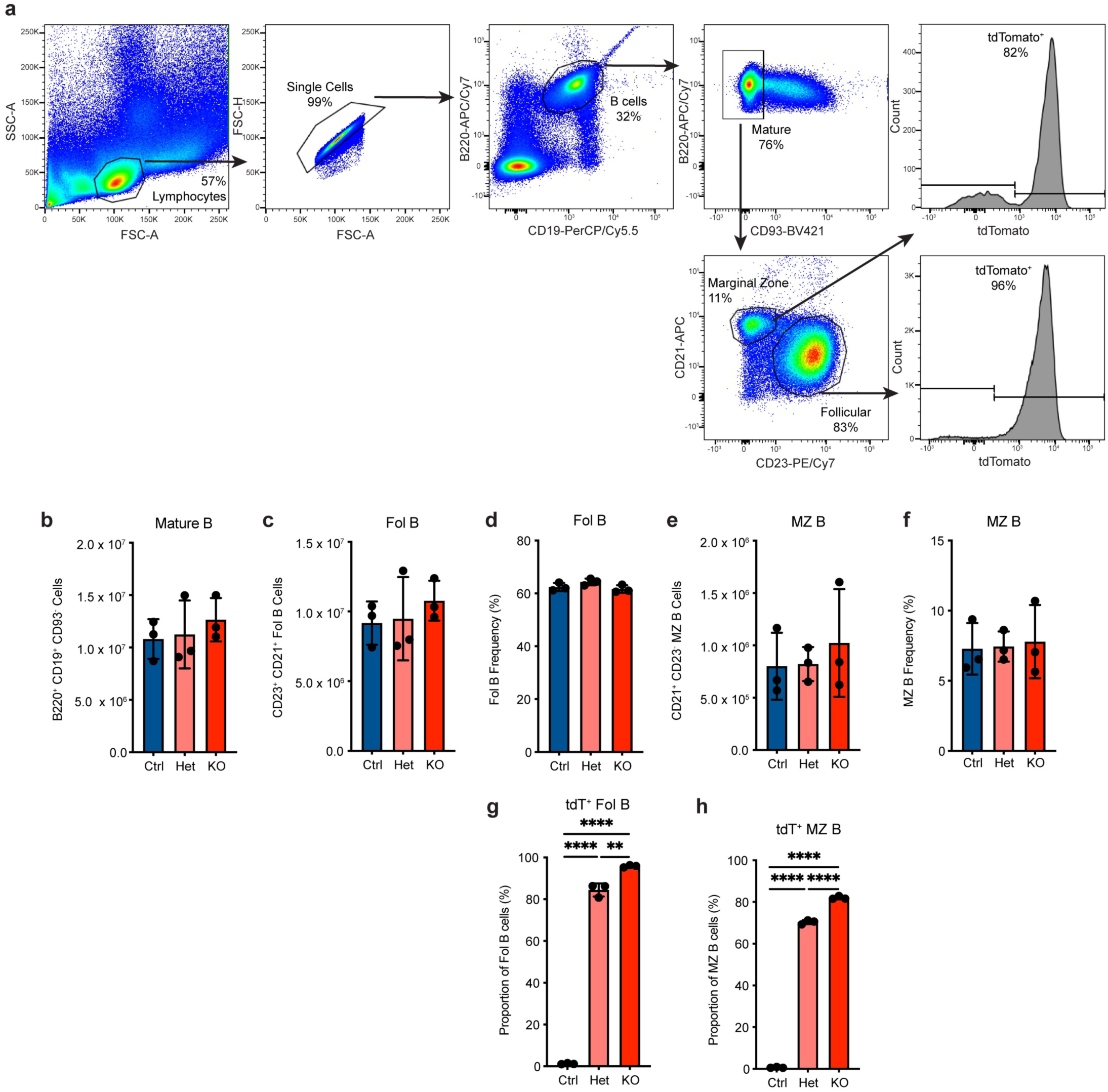
Normal Fol and MZ B cell numbers in *Brwd1*-KO^Fol^ mice. **(a**) Splenic Fol B cells were gated as B220^+^CD19^+^CD93^-^CD23^+^CD21^+^ cells by flow cytometry. Representative data from a *Cd23^Cre/wt^ Brwd1^fl/fl^* mouse is shown. **(b)** Number of B220^+^CD19^+^CD93^-^ mature B cells from *Brwd1^fl/fl^* (Ctrl), *Cd23^Cre/wt^ Brwd1^fl/wt^* (Het), and *Cd23^Cre/wt^ Brwd1^fl/fl^* (KO) mice (n = 3 mice per group). **(c)** Number of CD23^+^CD21^+^ Fol B cells. **(d)** Frequency of Fol B cells as a proportion of all B220^+^CD19^+^ cells. **(e)** Number of CD21^+^CD23^-^ MZ B cells. **(f)** Frequency of MZ B cells as a proportion of all B220^+^CD19^+^ cells. **(g)** Frequency of tdT^+^ Fol B cells. **(h)** Frequency of tdT^+^ MZ B cells. (\*\**p* < 0.01, \*\*\*\**p* < 0.0001, two-sided unpaired *t*-test, bar plots show mean ± standard deviation)

**Extended Data Figure 3.**
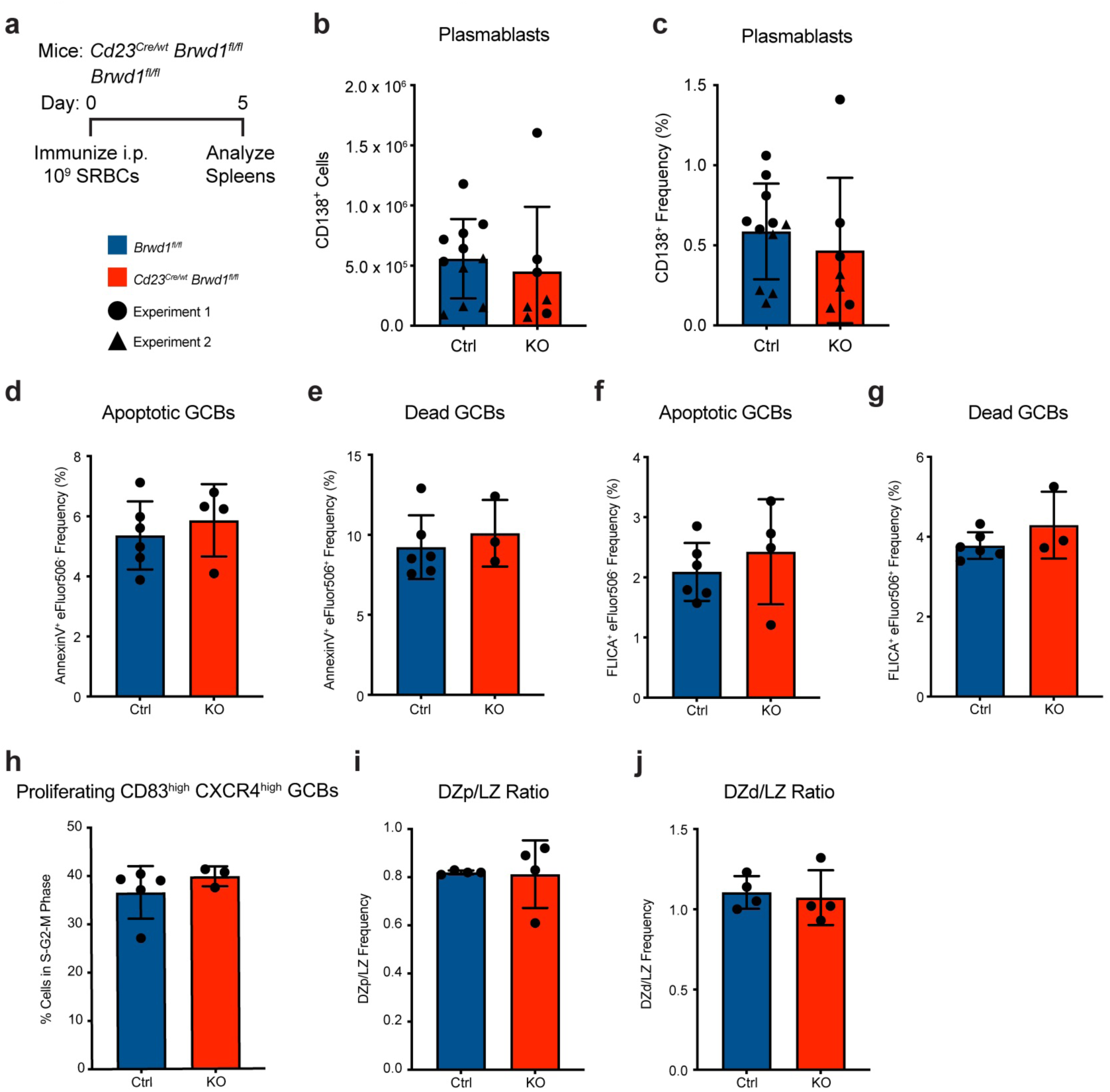
Normal plasmablasts and apoptosis in immunized *Brwd1*-KO^Fol^ mice. **(a)** *Cd23^Cre/wt^ Brwd1^fl/fl^* (KO) and *Brwd1^fl/fl^* (Ctrl) mice were immunized intraperitoneally (i.p.) with 10^9^ sheep red blood cells (SRBCs), and spleens were collected at day 5 post-immunization. **(b-c)** Total number (b) and frequency (c) of CD138^+^ plasmablasts (Ctrl n = 11 mice, KO n = 7 mice). **(d)** Frequency of apoptotic AnnexinV^+^eFluor506^-^ GC B cells. Data is gated on B220^+^CD95^+^GL7^+^ cells (Ctrl n = 6 mice, KO n = 4 mice). **(e)** Frequency of dead AnnexinV^+^eFluor506^+^ GC B cells. **(f)** Frequency of apoptotic FLICA^+^eFluor506^-^ GC B cells. **(g)** Frequency of dead FLICA^+^eFluor506^+^ GC B cells. **(h)** Frequency of DAPI^high^CD83^high^CXCR4^high^ DZp GC B cells (Ctrl n = 5 mice, KO n = 3 mice). **(i-j)** Ratios of DZp frequency to LZ frequency (i) and of DZd frequency to LZ frequency (j) (Ctrl n = 4 mice, KO n = 4 mice). (two-sided unpaired *t*-test, bar plots show mean ± standard deviation)

**Extended Data Figure 4.**
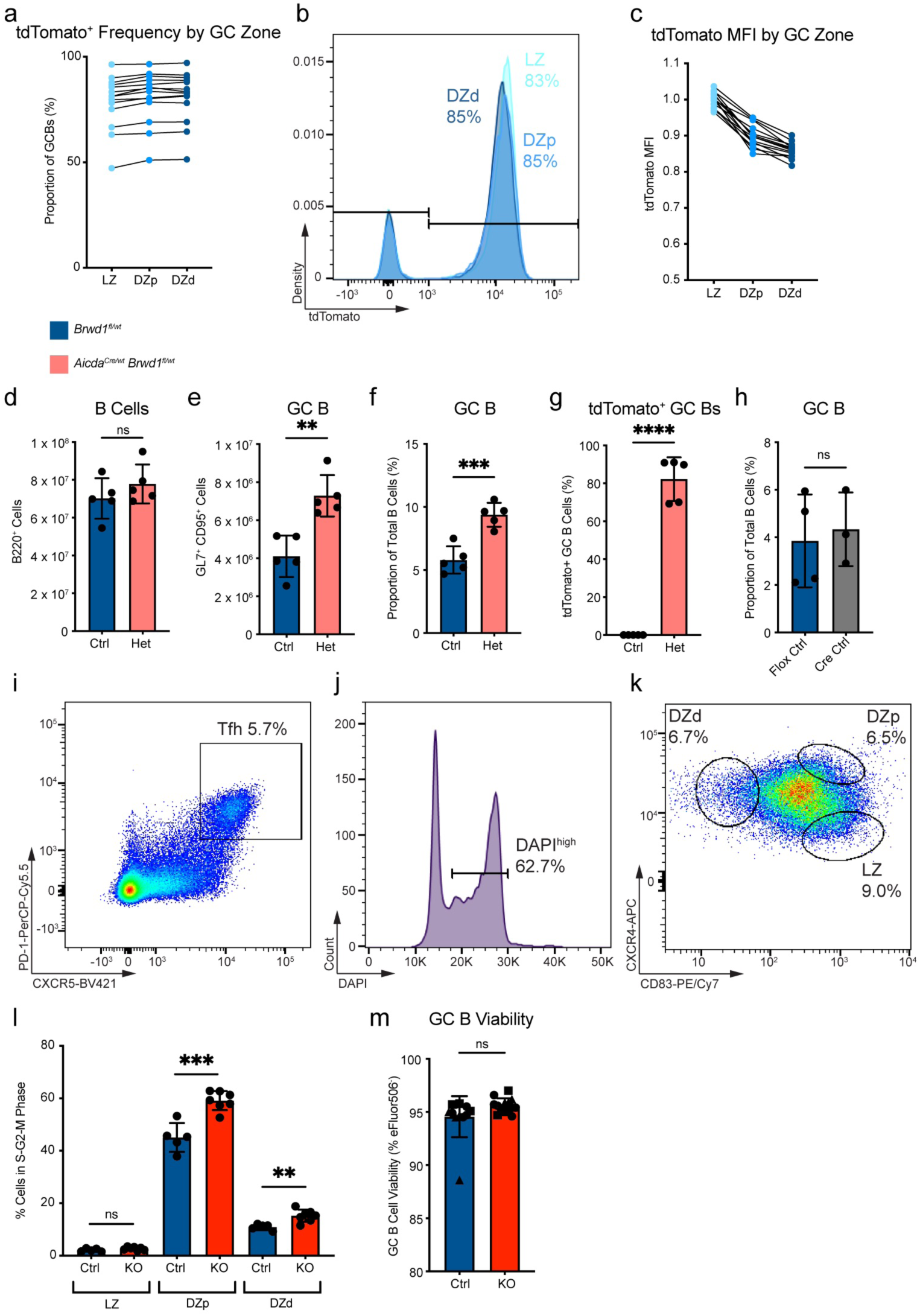
BRWD1 restrains DZp and DZd cell proliferation in the GC. **(a)** Frequency of tdT^+^ GC B cells across each GC subset of immunized *Aicda^Cre/wt^ Brwd1^fl/fl^* mice.Dots are connected for each mouse (n = 14 mice). **(b)** Representative flow plot of tdT fluorescence. **(c)** tdT median fluorescence intensity (MFI) across each GC subset. MFI was normalized due to differences in flow cytometry voltage settings between 3 independent experiments. **(d)** Total B cell number in *Aicda^Cre/wt^ Brwd1^fl/wt^* (Het) and *Brwd1^fl/wt^* (Ctrl) mice immunized with sheep red blood cells (SRBCs) and boosted at day 5. Splenic cells were analyzed at day 14 post-immunization (Ctrl n = 5 mice, Het n = 5 mice). **(e)** Total GC B cell number. **(f)** GC B cell frequency. **(g)** Proportion of GC B cells expressing tdT. **(h)** GC B cell frequency comparing *Brwd1^fl/fl^* (Flox Ctrl) and *Aicda^Cre/wt^* (Cre Ctrl) mice (Flox Ctrl n = 4 mice, Cre Ctrl n = 3 mice). **(i)** Representative flow plot of T follicular helper (Tfh) cell gating strategy. First gated on CD3^+^CD4^+^ cells. **(j)** Representative flow plot of DAPI^high^ gating strategy. First gated on B220^+^CD95^+^GL7^+^CD83^+^CXCR4^+^ DZp GC B cells. **(k)** Flow cytometry gating strategy of LZ, DZp, and DZd GC B cells. Previously gated on B220^+^GL7^+^CD95^+^ cells. **(l)** Frequency of proliferating GC B cells in the S, G2, or M phases of the cell cycle as measured by DAPI^high^ frequency (Ctrl n = 5 mice, KO n = 7 mice). **(m)** GC B cell viability measured by the frequency of the viability dye eFluor506^-^ cells (Ctrl n = 12 mice, KO n = 14 mice). (\*\**p* < 0.01, \*\*\**p* < 0.001, \*\*\*\**p* < 0.0001, two-sided unpaired *t*-test, bar plots show mean ± standard deviation)

**Extended Data Figure 5.**
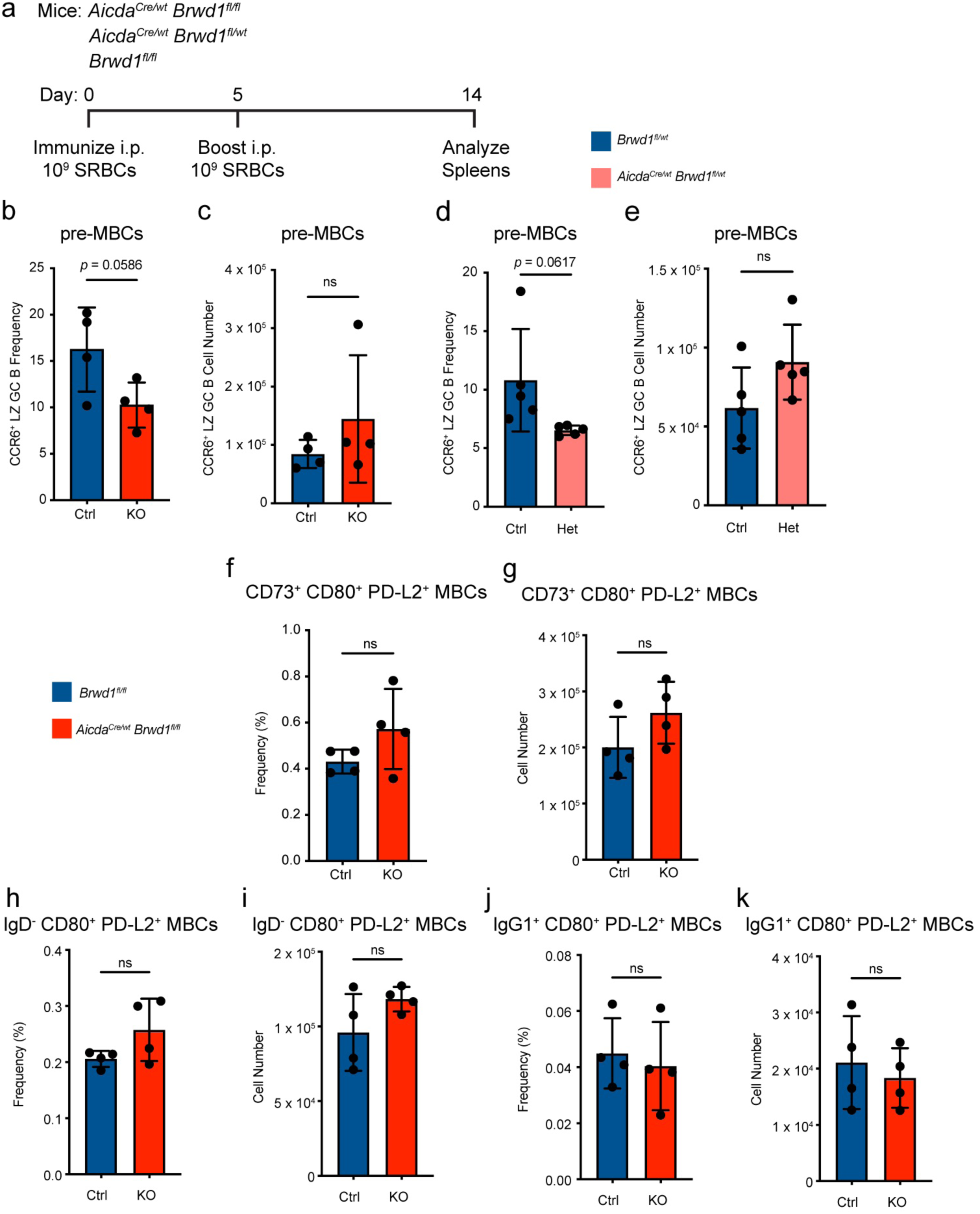
BRWD1 does not regulate post-GC MBCs or PCs. **(a)** *Aicda^Cre/wt^ Brwd1^fl/fl^* (KO) and *Brwd1^fl/fl^* (Ctrl) mice were immunized with SRBCs i.p. and boosted at day 5. Splenic cells were analyzed at day 14 post-immunization. **(b-c)** Frequency (b) and number (c) of CCR6^+^CD83^high^ pre-MBCs (Ctrl n = 4 mice, KO n = 4 mice). **(d-e)** Frequency (d) and number (e) of CCR6^+^CD83^high^ pre-MBCs in *Aicda^Cre/wt^ Brwd1^fl/wt^* (Het) and *Brwd1^fl/wt^* (Ctrl) mice (Ctrl n = 5 mice, Het n = 5 mice). **(f-k)** Frequency or number of different CD80^+^PD-L2^+^ MBC populations either gating first on CD73^+^ cells (f-g), IgD^-^ cells (h-i), or IgG1^+^ cells (j-k). Frequencies are calculated as a proportion of B220^+^CD38^+^GL7^-^ cells (Ctrl n = 4 mice, KO n = 4 mice). (two-sided unpaired *t*-test, bar plots show mean ± standard deviation)

**Extended Data Figure 6.**
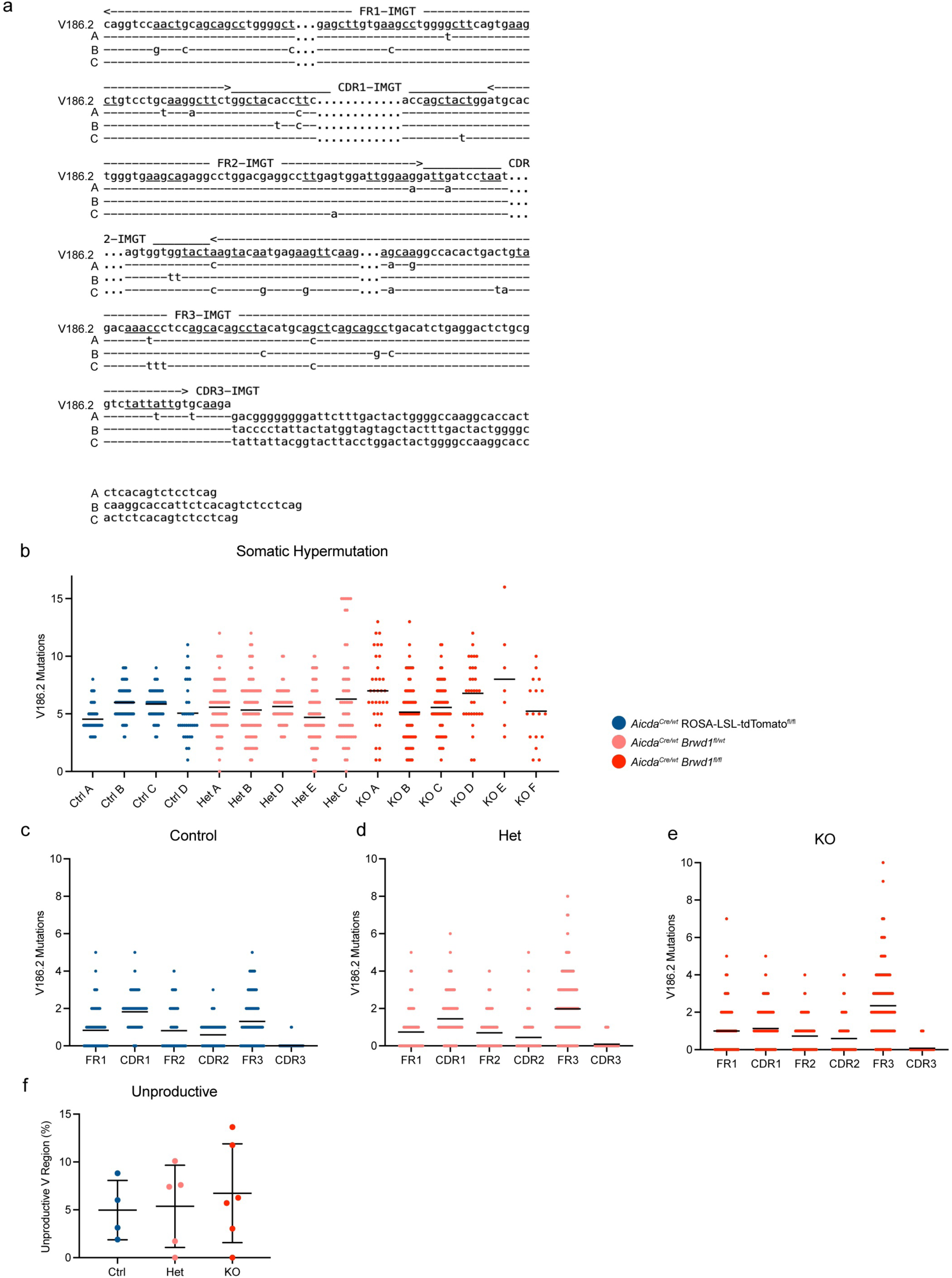
*Brwd1*^-/-^ GC B cells manifest wide distributions in SHM frequencies. **(a)** Example distribution of SHM in three representative *Aicda^Cre/wt^ Brwd1^fl/fl^* (KO) tdT^+^ GC B cells with a relatively high level of mutations. The V_H_186.2 nucleotide sequence is shown above. **(b)** Frequency of mutations within V_H_186.2 in GC B cells per mouse. Lines show mean (Ctrl n = 4 mice, Het n = 5 mice, KO n = 6 mice). **(c-e)** Distribution of V_H_186.2 mutations within framework regions (FR) and complementarity-determining regions (CDR) for *Aicda^Cre/wt^* ROSA26-LSL-tdTomato^fl/fl^ (Ctrl) GC B cells (c), *Aicda^Cre/wt^ Brwd1^fl/wt^* (Het) GC B cells (d), and *Aicda^Cre/wt^ Brwd1^fl/fl^* (KO) GC B cells (e). Mutations in CDR3 introduced by junctional diversity were not counted. **(f)** Frequency of clones with an unproductive V region. Lines show mean ± standard deviation.

**Extended Data Figure 7.**
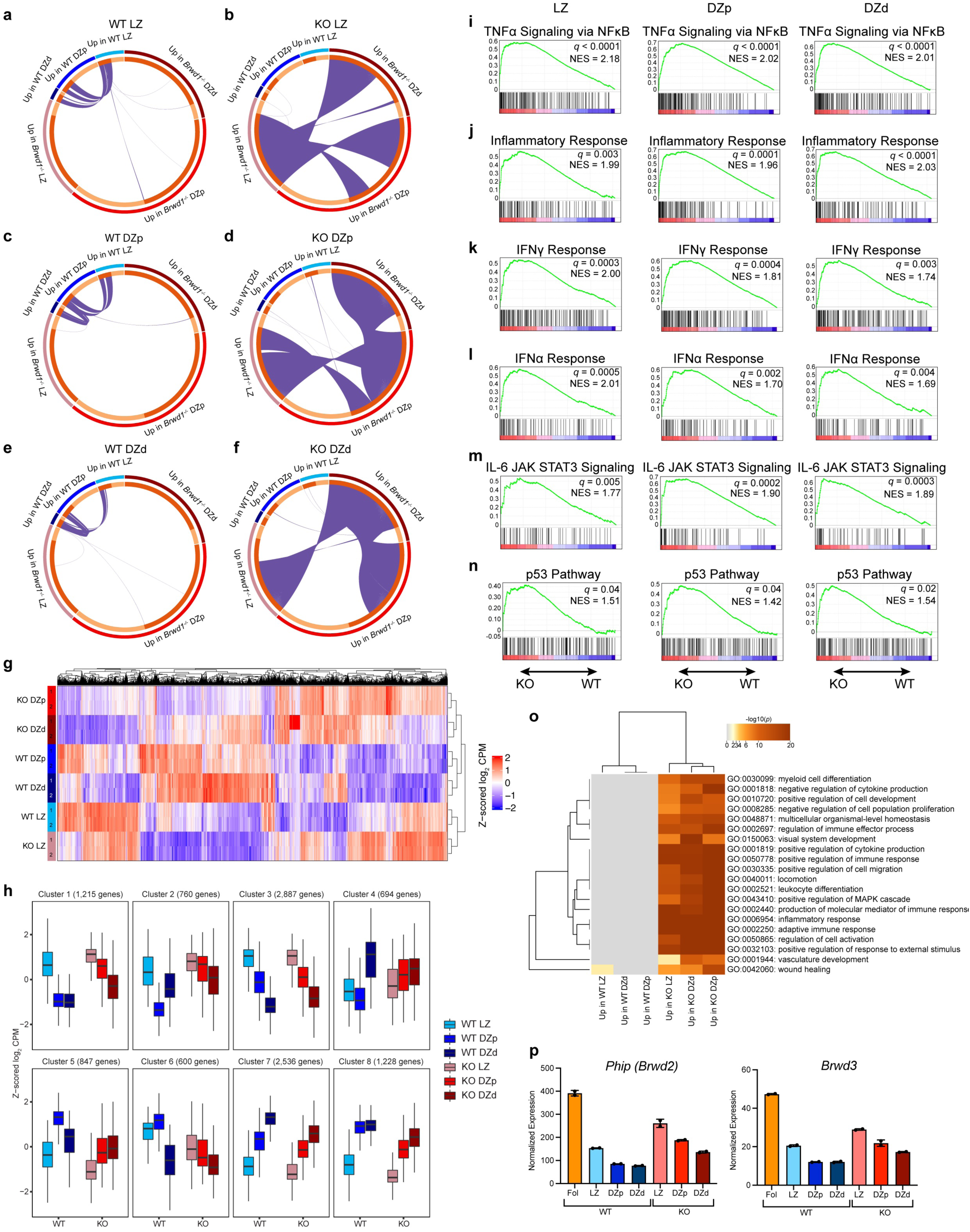
BRWD1 maintains GC B cell subset transcriptional identity. **(a-f)** Differentially expressed genes between *Brwd1^-/-^* and WT GC B cells from the LZ, DZp, or DZd. Purple lines show genes shared between groups. Dark orange signifies genes in the group that are shared with another group. Each plot shows shared genes from the perspective of the WT LZ (a), KO LZ (b), WT DZp (c), KO DZp (d), WT DZd (e), or KO DZd (f). n = 2 per cell type. Each n represents cells pooled from 20 mice. **(g)** Heatmap of 11,998 differentially expressed genes (one-way ANOVA, *q* < 0.05) between KO and WT GC B cells from the LZ, DZp, or DZd. Z-scored log_2_ counts per million (CPM) are plotted. **(h)** Gene expression of WT and KO GC subsets at clusters generated by unsupervised *K*-means clustering of WT GC subsets, performed previously^14^. Z-scored log_2_ CPM are plotted. Box limits show interquartile range (IQR), and center lines show median. Maximum and minimum values (whiskers) are defined as Q3 + 1.5 x IQR and Q1 – 1.5 x IQR, respectively. **(i-n)** Gene set enrichment analyses for the TNFα signaling via NFκB (i), inflammatory response (j), IFNγ response (k), IFNα response (l), IL-6 JAK-STAT3 signaling (m), and p53 (n) pathways for each GC subpopulation. **(o)** Gene Ontology (GO) pathways enriched in differentially expressed genes between *Brwd1^-/-^* and WT cells from the LZ, DZp, or DZd. The top 20 pathways by *p* value are listed. **(p)** RNA-seq expression of *Phip* (also known as *Brwd2*) and *Brwd3*. Bar plots show mean ± standard deviation.

**Supplementary Table 1.**
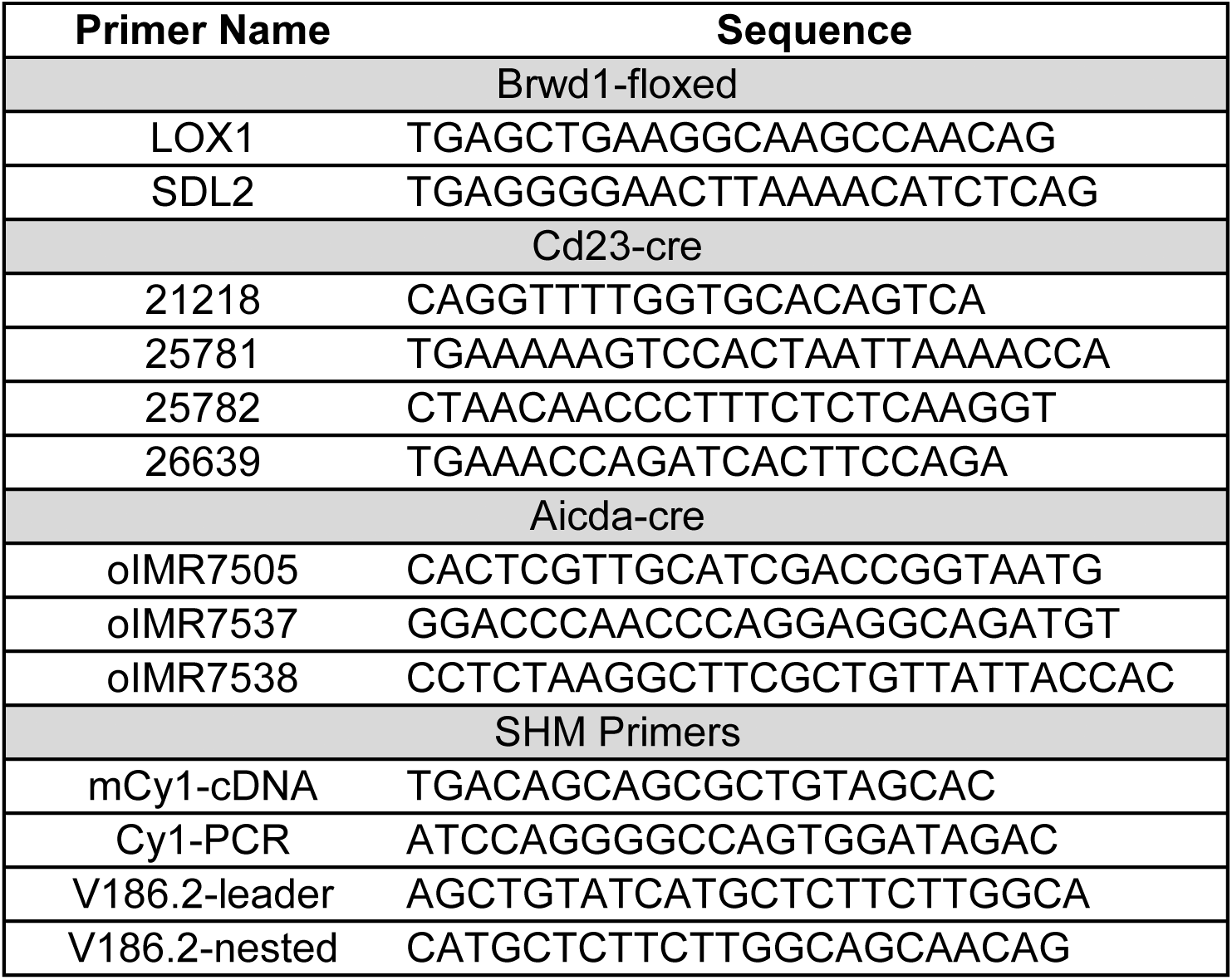
List of Primers.

## REFERENCES

1. Cyster, J. G. & Allen, C. D. C. B cell responses: cell interaction dynamics and decisions. Cell 177, 524–540 (2019).

2. Okada, T. et al. Antigen-engaged B cells undergo chemotaxis toward the T zone and form motile conjugates with helper T cells. PLoS Biol. 3, e150 (2005).

3. Kerfoot, S. M. et al. Germinal center B cell and T follicular helper cell development initiates in the interfollicular zone. Immunity 34, 947–960 (2011).

4. Kitano, M. et al. Bcl6 protein expression shapes pre-germinal center B cell dynamics and follicular helper T cell heterogeneity. Immunity 34, 961–972 (2011).

5. Zhang, T. et al. Germinal center B cell development has distinctly regulated stages completed by disengagement from T cell help. Elife 6, e19552 (2017).

6. Robinson, M. J. et al. The amount of BCL6 in B cells shortly after antigen engagement determines their representation in subsequent germinal centers. Cell Reports 30, 1530–1541.e4 (2020).

7. Huang, C. et al. The BCL6 RD2 domain governs commitment of activated B cells to form germinal centers. Cell Reports 8, 1497–1508 (2014).

8. Silva, N. S. D. & Klein, U. Dynamics of B cells in germinal centres. Nat Rev Immunol 15, 137– 148 (2015).

9. Papin, A., Cesarman, E. & Melnick, A. 3D chromosomal architecture in germinal center B cells and its alterations in lymphomagenesis. Curr Opin Genet Dev 74, 101915 (2022).

10. Vilarrasa-Blasi, R. et al. Dynamics of genome architecture and chromatin function during human B cell differentiation and neoplastic transformation. Nat Commun 12, 651 (2021).

11. Bunting, K. L. et al. Multi-tiered reorganization of the genome during B cell affinity maturation anchored by a germinal center-specific locus control region. Immunity 45, 497–512 (2016).

12. Doane, A. S. et al. OCT2 pre-positioning facilitates cell fate transition and chromatin architecture changes in humoral immunity. Nat Immunol 22, 1327–1340 (2021).

13. Roco, J. A. et al. Class-switch recombination occurs infrequently in germinal centers. Immunity 51, 337–350.e7 (2019).

14. Kennedy, D. E. et al. Novel specialized cell state and spatial compartments within the germinal center. Nat Immunol 21, 660–670 (2020).

15. Wright, N. E., Mandal, M. & Clark, M. R. Molecular mechanisms insulating proliferation from genotoxic stress in B lymphocytes. Trends Immunol. 44, 668–677 (2023).

16. Kennedy, D. E. & Clark, M. R. Compartments and connections within the germinal center. Front Immunol 12, 659151 (2021).

17. Victora, G. D. et al. Germinal center dynamics revealed by multiphoton microscopy with a photoactivatable fluorescent reporter. Cell 143, 592–605 (2010).

18. Allen, C. D. C., Okada, T., Tang, H. L. & Cyster, J. G. Imaging of germinal center selection events during affinity maturation. Science 315, 528–531 (2007).

19. Liu, D. et al. T–B-cell entanglement and ICOSL-driven feed-forward regulation of germinal centre reaction. Nature 517, 214–218 (2015).

20. Gitlin, A. D., Shulman, Z. & Nussenzweig, M. C. Clonal selection in the germinal centre by regulated proliferation and hypermutation. Nature 509, 637–640 (2014).

21. Shulman, Z. et al. Dynamic signaling by T follicular helper cells during germinal center B cell selection. Science 345, 1058–1062 (2014).

22. Ise, W. et al. T follicular helper cell-germinal center B cell interaction strength regulates entry into plasma cell or recycling germinal center cell fate. Immunity 48, 702–715.e4 (2018).

23. Ersching, J. et al. Germinal center selection and affinity maturation require dynamic regulation of mTORC1 kinase. Immunity 46, 1045–1058.e6 (2017).

24. Luo, W., Weisel, F. & Shlomchik, M. J. B cell receptor and CD40 signaling are rewired for synergistic induction of the c-Myc transcription factor in germinal center B cells. Immunity 48, 313–326.e5 (2018).

25. Chen, S. T., Oliveira, T. Y., Gazumyan, A., Cipolla, M. & Nussenzweig, M. C. B cell receptor signaling in germinal centers prolongs survival and primes B cells for selection. Immunity 56, 547–561.e7 (2023).

26. Duan, L. et al. Follicular dendritic cells restrict interleukin-4 availability in germinal centers and foster memory B cell generation. Immunity 54, 2256–2272.e6 (2021).

27. Long, Z., Phillips, B., Radtke, D., Meyer-Hermann, M. & Bannard, O. Competition for refueling rather than cyclic reentry initiation evident in germinal centers. Sci Immunol 7, eabm0775 (2022).

28. Clark, M. R., Mandal, M., Ochiai, K. & Singh, H. Orchestrating B cell lymphopoiesis through interplay of IL-7 receptor and pre-B cell receptor signalling. Nat Rev Immunol 14, 69–80 (2014).

29. McLean, K. C. & Mandal, M. It takes three receptors to raise a B cell. Trends Immunol 41, 629–642 (2020).

30. Mandal, M. et al. Histone reader BRWD1 targets and restricts recombination to the Igk locus. Nat Immunol 16, 1094–1103 (2015).

31. Mandal, M. et al. BRWD1 orchestrates epigenetic landscape of late B lymphopoiesis. Nat Commun 9, 3888 (2018).

32. Mandal, M. et al. BRWD1 orchestrates small pre-B cell chromatin topology by converting static to dynamic cohesin. Nat. Immunol. 25, 129–141 (2024).

33. Kwon, K. et al. Instructive role of the transcription factor E2A in early B lymphopoiesis and germinal center B cell development. Immunity 28, 751–762 (2008).

34. Weisel, F. J., Zuccarino-Catania, G. V., Chikina, M. & Shlomchik, M. J. A temporal switch in the germinal center determines differential output of memory B and plasma cells. Immunity 44, 116–130 (2016).

35. Merkenschlager, J. et al. Dynamic regulation of TFH selection during the germinal centre reaction. Nature 591, 458–463 (2021).

36. Zhang, Y., Zhang, X., Dai, H.-Q., Hu, H. & Alt, F. W. The role of chromatin loop extrusion in antibody diversification. Nat Rev Immunol 22, 550–566 (2022).

37. Zhang, X. et al. Fundamental roles of chromatin loop extrusion in antibody class switching. Nature 575, 385–389 (2019).

38. Wuerffel, R. et al. S-S synapsis during class switch recombination is promoted by distantly located transcriptional elements and activation-induced deaminase. Immunity 27, 711–722 (2007).

39. Song, S. et al. OBF1 and Oct factors control the germinal center transcriptional program. Blood 137, 2920–2934 (2021).

40. Hodson, D. J. et al. Regulation of normal B-cell differentiation and malignant B-cell survival by OCT2. Proc National Acad Sci 113, E2039–E2046 (2016).

41. Sunshine, A. et al. Ets1 controls the development of B cell autoimmune responses in a cell-intrinsic manner. ImmunoHorizons 3, 331–340 (2019).

42. Luo, W. et al. A balance between B cell receptor and inhibitory receptor signaling controls plasma cell differentiation by maintaining optimal Ets1 levels. J. Immunol. 193, 909–920 (2014).

43. Willis, S. N. et al. Environmental sensing by mature B cells is controlled by the transcription factors PU.1 and SpiB. Nat Commun 8, 1426 (2017).

44. Bonetti, P. et al. Deregulation of ETS1 and FLI1 contributes to the pathogenesis of diffuse large B-cell lymphoma. Blood 122, 2233–2241 (2013).

45. Xu, H. et al. Regulation of bifurcating B cell trajectories by mutual antagonism between transcription factors IRF4 and IRF8. Nat Immunol 16, 1274–1281 (2015).

46. Ochiai, K. et al. Transcriptional regulation of germinal center B and plasma cell fates by dynamical control of IRF4. Immunity 38, 918–929 (2013).

47. Crouch, E. E. et al. Regulation of AID expression in the immune response. J Exp Medicine 204, 1145–1156 (2007).

48. Suan, D. et al. CCR6 defines memory B cell precursors in mouse and human germinal centers, revealing light-zone location and predominant low antigen affinity. Immunity 47, 1142–1153.e4 (2017).

49. Laidlaw, B. J. et al. The Eph-related tyrosine kinase ligand Ephrin-B1 marks germinal center and memory precursor B cells. J Exp Medicine 214, 639–649 (2017).

50. Inoue, T. et al. Exit from germinal center to become quiescent memory B cells depends on metabolic reprograming and provision of a survival signal. J Exp Med 218, e20200866 (2020).

51. Viant, C. et al. Germinal center–dependent and –independent memory B cells produced throughout the immune response. J Exp Med 218, e20202489 (2021).

52. Callahan, D. et al. Memory B cell subsets have divergent developmental origins that are coupled to distinct imprinted epigenetic states. Nat. Immunol. 25, 562–575 (2024).

53. Zuccarino-Catania, G. V. et al. CD80 and PD-L2 define functionally distinct memory B cell subsets that are independent of antibody isotype. Nat Immunol 15, 631–637 (2014).

54. Kaji, T. et al. Distinct cellular pathways select germline-encoded and somatically mutated antibodies into immunological memory. J Exp Med 209, 2079–2097 (2012).

55. Taylor, J. J., Pape, K. A. & Jenkins, M. K. A germinal center–independent pathway generates unswitched memory B cells early in the primary response. J Exp Medicine 209, 597–606 (2012).

56. Conter, L. J., Song, E., Shlomchik, M. J. & Tomayko, M. M. CD73 expression is dynamically regulated in the germinal center and bone marrow plasma cells are diminished in its absence. PloS One 9, e92009 (2014).

57. Heise, N. & Klein, U. Somatic hypermutation and affinity maturation analysis using the 4-hydroxy-3-nitrophenyl-acetyl (NP) system. Methods Mol Biology 1623, 191–208 (2017).

58. Finkin, S., Hartweger, H., Oliveira, T. Y., Kara, E. E. & Nussenzweig, M. C. Protein amounts of the MYC transcription factor determine germinal center B cell division capacity. Immunity 51, 324–336.e5 (2019).

59. Pae, J. et al. Cyclin D3 drives inertial cell cycling in dark zone germinal center B cells. J Exp Med 218, e20201699 (2020).

60. Weiser, A. A. et al. Affinity maturation of B cells involves not only a few but a whole spectrum of relevant mutations. Int Immunol 23, 345–356 (2011).

61. Burnett, D. L. et al. Germinal center antibody mutation trajectories are determined by rapid self/foreign discrimination. Science 360, 223–226 (2018).

62. Mayer, C. T. et al. The microanatomic segregation of selection by apoptosis in the germinal center. Science 358, eaao2602 (2017).

63. Butt, D. et al. FAS inactivation releases unconventional germinal center B cells that escape antigen control and drive IgE and autoantibody production. Immunity 42, 890–902 (2015).

64. Chan, T. D. et al. Elimination of germinal-center-derived self-reactive B cells is governed by the location and concentration of self-antigen. Immunity 37, 893–904 (2012).

65. Okoreeh, M. K., et al. Asymmetrical forward and reverse developmental trajectories determine molecular programs of B cell antigen receptor editing. Sci Immunol 7, eabm1664 (2022).

66. Matz, H. & Dooley, H. 450 million years in the making: mapping the evolutionary foundations of germinal centers. Front. Immunol. 14, 1245704 (2023).

67. Dooley, H. & Flajnik, M. F. Shark immunity bites back: affinity maturation and memory response in the nurse shark, Ginglymostoma cirratum. Eur. J. Immunol. 35, 936–945 (2005).

68. Castro, C. D., Ohta, Y., Dooley, H. & Flajnik, M. F. Noncoordinate expression of J-chain and Blimp-1 define nurse shark plasma cell populations during ontogeny. Eur. J. Immunol. 43, 3061– 3075 (2013).

69. Dooley, H., Stanfield, R. L., Brady, R. A. & Flajnik, M. F. First molecular and biochemical analysis of in vivo affinity maturation in an ectothermic vertebrate. Proc. Natl. Acad. Sci. 103, 1846–1851 (2006).

70. Matz, H. et al. Organized B cell sites in cartilaginous fishes reveal the evolutionary foundation of germinal centers. Cell Rep. 42, 112664 (2023).

71. Davidson, I. F. & Peters, J.-M. Genome folding through loop extrusion by SMC complexes. Nat Rev Mol Cell Bio 22, 445–464 (2021).

72. Busslinger, G. A. et al. Cohesin is positioned in mammalian genomes by transcription, CTCF and Wapl. Nature 544, 503–507 (2017).

73. Thiecke, M. J. et al. Cohesin-dependent and -independent mechanisms mediate chromosomal contacts between promoters and enhancers. Cell Rep. 32, 107929 (2020).

74. Khattabi, L. E. et al. A pliable Mediator acts as a functional rather than an architectural bridge between promoters and enhancers. Cell 178, 1145–1158.e20 (2019).

75. Cato, M. H., Yau, I. W. & Rickert, R. C. Magnetic-based purification of untouched mouse germinal center B cells for ex vivo manipulation and biochemical analysis. Nat Protoc 6, 953– 960 (2011).

76. Schindelin, J., et al. Fiji: an open-source platform for biological-image analysis. Nat. Methods 9, 676–682 (2012).

77. Walt, S. van der et al. scikit-image: image processing in Python. PeerJ 2, e453 (2014).

78. Brochet, X., Lefranc, M.-P. & Giudicelli, V. IMGT/V-QUEST: the highly customized and integrated system for IG and TR standardized V-J and V-D-J sequence analysis. Nucleic Acids Res. 36, W503–W508 (2008).

79. Dobin, A. et al. STAR: ultrafast universal RNA-seq aligner. Bioinformatics 29, 15–21 (2013).

80. Liao, Y., Smyth, G. K. & Shi, W. featureCounts: an efficient general purpose program for assigning sequence reads to genomic features. Bioinformatics 30, 923–930 (2014).

81. Li, H. Aligning sequence reads, clone sequences and assembly contigs with BWA-MEM. arXiv https://arxiv.org/abs/1303.3997 (2013).

82. Quinlan, A. R. & Hall, I. M. BEDTools: a flexible suite of utilities for comparing genomic features. Bioinformatics 26, 841–842 (2010).

83. Zhang, Y. et al. Model-based analysis of ChIP-seq (MACS). Genome Biol. 9, R137 (2008).

84. Robinson, M. D., McCarthy, D. J. & Smyth, G. K. edgeR: a Bioconductor package for differential expression analysis of digital gene expression data. Bioinformatics 26, 139–140 (2010).

85. Freese, N. H., Norris, D. C. & Loraine, A. E. Integrated genome browser: visual analytics platform for genomics. Bioinformatics 32, 2089–2095 (2016).

86. Gu, Z., Eils, R. & Schlesner, M. Complex heatmaps reveal patterns and correlations in multidimensional genomic data. Bioinformatics 32, 2847–2849 (2016).

87. Gu, Z., Gu, L., Eils, R., Schlesner, M. & Brors, B. circlize implements and enhances circular visualization in R. Bioinformatics 30, 2811–2812 (2014).

88. Zhou, Y. et al. Metascape provides a biologist-oriented resource for the analysis of systems-level datasets. Nat. Commun. 10, 1523 (2019).

89. Heinz, S. et al. Simple combinations of lineage-determining transcription factors prime cis-regulatory elements required for macrophage and B cell identities. Mol Cell 38, 576–589 (2010).

90. Subramanian, A. et al. Gene set enrichment analysis: A knowledge-based approach for interpreting genome-wide expression profiles. Proc. Natl. Acad. Sci. 102, 15545–15550 (2005).

91. Liberzon, A. et al. The molecular signatures database hallmark gene set collection. Cell Syst. 1, 417–425 (2015).

92. Robinson, J. T. et al. Juicebox.js provides a cloud-based visualization system for Hi-C data. Cell Syst. 6, 256–258.e1 (2018).

93. Stansfield, J. C., Cresswell, K. G., Vladimirov, V. I. & Dozmorov, M. G. HiCcompare: an R-package for joint normalization and comparison of HI-C datasets. BMC Bioinform. 19, 279 (2018).

94. Phanstiel, D. H., Boyle, A. P., Araya, C. L. & Snyder, M. P. Sushi.R: flexible, quantitative and integrative genomic visualizations for publication-quality multi-panel figures. Bioinformatics 30, 2808–2810 (2014).

